# Unveiling IRF4-steered regulation of context-dependent effector programs in Th17 and Treg cells

**DOI:** 10.1101/2023.09.14.557376

**Authors:** Anna Gabele, Maximilian Sprang, Mert Cihan, Sarah Dietzen, Matthias Klein, Gregory Harms, Tanja Ziesmann, Katrin Pape, Beatrice Wasser, David Gomez-Zepeda, Kathrin Braband, Michael Delacher, Niels Lemmermann, Stefan Bittner, Miguel A. Andrade-Navarro, Stefan Tenzer, Tobias Bopp, Ute Distler

**Affiliations:** Institute of Immunology, University Medical Center of the Johannes Gutenberg University Mainz, Mainz, Germany; Research Center for Immunotherapy (FZI), University Medical Center of the Johannes Gutenberg University Mainz, Mainz, Germany; Faculty of Biology, Johannes Gutenberg University Mainz, Mainz, Germany; Cell Biology Unit, University Medical Center of the Johannes Gutenberg University Mainz, Mainz, Germany; Department of Neurology, University Medical Center of the Johannes Gutenberg University Mainz, Mainz, Germany; Helmholtz Institute for Translational Oncology (HI-TRON), Mainz, Germany; Deutsches Krebsforschungszentrum (DKFZ), Heidelberg, Germany; Institute for Virology, University Medical Center of the Johannes Gutenberg University Mainz, Mainz, Germany; Institute of Virology, Medical Faculty, University Bonn, Germany; Focus Program Translational Neuroscience (FTN), University Medical Center of the Johannes Gutenberg University Mainz, Mainz, Germany

## Abstract

The transcription factor interferon regulatory factor 4 (IRF4) is crucial for the differentiation and fate determination of pro-inflammatory T helper (Th)17 and the functionally opposing group of immunomodulatory regulatory T (Treg) cells. However, molecular mechanisms of how IRF4 steers diverse transcriptional programs in Th17 and Treg cells are far from being definitive. To unveil IRF4-driven lineage determination in Th17 and Treg cells, we integrated data derived from affinity-purification and full mass spectrometry-based proteome analysis with chromatin immune precipitation sequencing (ChIP-Seq). This allowed the characterization of subtype-specific molecular programs and the identification of novel, previously unknown IRF4 interactors in the Th17/Treg context, such as RORγt, AHR, IRF8, BACH2, SATB1, and FLI1. Moreover, our data reveal that most of these transcription factors are recruited to IRF composite elements for the regulation of cell type-specific transcriptional programs providing a valuable resource for studying IRF4-mediated gene regulatory programs in pro- and anti-inflammatory immune responses.

## INTRODUCTION

The transcription factor (TF) interferon regulatory factor 4 (IRF4) is one of nine members in the mammalian family of interferon regulatory factors (IRFs). IRFs steer different aspects of innate and adaptive immune responses as they play a crucial role for the induction of type I interferons (IFN-α and -β) and are required for the development and function of different immune cells^1–3^. Unlike other members of the IRF-family, IRF4 is not regulated by interferons but induced by T cell receptor (TCR) signaling as well as by Toll-like and tumor necrosis factor receptors^2^. IRF4 expression is restricted to immune cells, such as T or B lymphocytes, dendritic cells, and macrophages. As IRF4 acts as one of the key regulators in the differentiation of those cell types, dysregulation of IRF4 is often associated with diseases like cancer and autoimmune disorders as, for example, multiple sclerosis. Pro-inflammatory Th17 cells, which have been implicated in the initiation and progression of autoimmune diseases, heavily rely on the expression of IRF4 during their development as demonstrated for the first time in 2007 by Brüstle *et al*.^3,4^. Immunoregulatory Treg cells, which contribute to the maintenance of peripheral tolerance by preventing auto-aggressive immune responses, do not directly rely on IRF4 for their generation^5^. However, the TF is essential for the context- and tissue-specific effector functions of this T cell type. Therefore, IRF4 is a crucial player, which contributes to the important and delicate equilibrium between pro- and anti-inflammatory immune responses as it is essential for the development of fully functional effector Th17 and Treg cells^2,5,6^.

Despite its central role in Th17 and Treg cell lineage determination and cellular plasticity, certain molecular mechanisms of IRF4-mediated gene expression in these cell types are still not fully understood. Studies performed over the past decade have revealed IRF4 target genes steering murine and human Th17 and Treg effector cell differentiation and function. For example, it could be demonstrated, that IRF4 directly binds at key lineage-associated loci in Th17 cells (*Il17a, Il17f, Il12rb1, Il1r1, Rorc*)^7,8^ and lack of IRF4 leads to a decreased expression of RORγt, the lineage-defining TF for Th17 cells^3^. In Treg cells, IRF4, in concert with BATF3, regulates the expression of the lineage-specific TF FOXP3 and hence controls (i)Treg induction^9^. Despite its repressive function on FOXP3 expression, IRF4 acts as a downstream target of FOXP3 and is required for the development of fully functional Treg cells and their immunomodulatory activity in a cellular context^6,9^. In a complex, IRF4 and FOXP3 control the expression of gene loci associated with Treg cell suppressive properties, e.g., *Il10* and *Icos*^10,11^. Moreover, it could be demonstrated that IRF4 directly controls a molecular program associated with increased immunosuppression in human tumors and abundance of intratumoral IRF4^+^ Treg cells correlates with poor prognosis in cancer patients^12^. Although respective studies have elucidated IRF4-mediated transcriptional programs and greatly contributed to our understanding of IRF4-steered molecular mechanisms in Th17 and Treg cells, major focus was set on target genes transcribed by IRF4, often at early time points during cellular differentiation (e.g., after 48 hours)^7,9,13^. However, gene expression is controlled by multiple molecular systems but only a very limited number of studies so far integrated information derived from IRF4 ChIP-Seq data with other ‘omics’ data at a global level in an unbiased manner (e.g. with transcriptomics^7^ or proteomics analysis) or shed, for example, light on proteins that regulate gene transcription in concert with IRF4 in a T cell subset-specific manner. Here, our current knowledge is mainly set on a few distinct binding partners^8,10,11,14^ and the question of how IRF4 interacting proteins steer IRF4-mediated target gene transcription in fully differentiated Th cell subtypes contributing to the cellular plasticity between Treg and Th17 cells remains unresolved to date^5^.

Affinity purification (AP) methods, such as the streptavidin capture of (*in vivo*) biotinylated proteins, coupled with mass spectrometry (MS) have emerged as powerful tools for the identification of novel protein-protein interactions ^15^. In 2005, Driegen *et al*. introduced a transgenic mouse strain ubiquitously expressing the bacterial BirA protein biotin ligase from the ROSA26 locus (ROSA26^BirA^ mice)^16^. BirA specifically biotinylates a short peptide sequence, also referred to as Avi-tag. ROSA26^BirA^ mice can be crossed with any transgenic mouse expressing an Avi-tagged protein to efficiently biotinylate the protein of interest for the purpose of protein (complex) purification. Beside the characterization of protein-protein interaction partners, this approach also enables the identification of target genes regulated by the TF (complex) of interest applying streptavidin-mediated chromatin precipitation coupled with deep sequencing (Bio-ChIP-Seq)^17,18^. In the present study, we integrate both strategies, AP-MS and Bio-ChIP-Seq, with MS-based proteome analyses to reveal IRF4-driven lineage determination in Treg and Th17 cells. Our data unraveled a complex and T cell subset-specific network of IRF-steered transcriptional regulation of T cell differentiation and plasticity.

## RESULTS

### IRF4 transcriptionally regulates distinct effector functions in Th17 and Treg cells

Comparative analyses of IRF4 expression in Th17 and Treg cells revealed that in both cell types IRF4 is expressed at a similar protein level (Fig. 1a). To better understand the impact of IRF4-mediated gene expression and its role for cellular plasticity in Th17 and Treg cells, we analyzed the full proteome of both cell types after *ex vivo* differentiation of naïve CD4^+^ T cells derived from conditional IRF4-deficient (*Irf4^−/−^*) and littermate control (WT) mice (Fig. 1b).

**Fig. 1.**
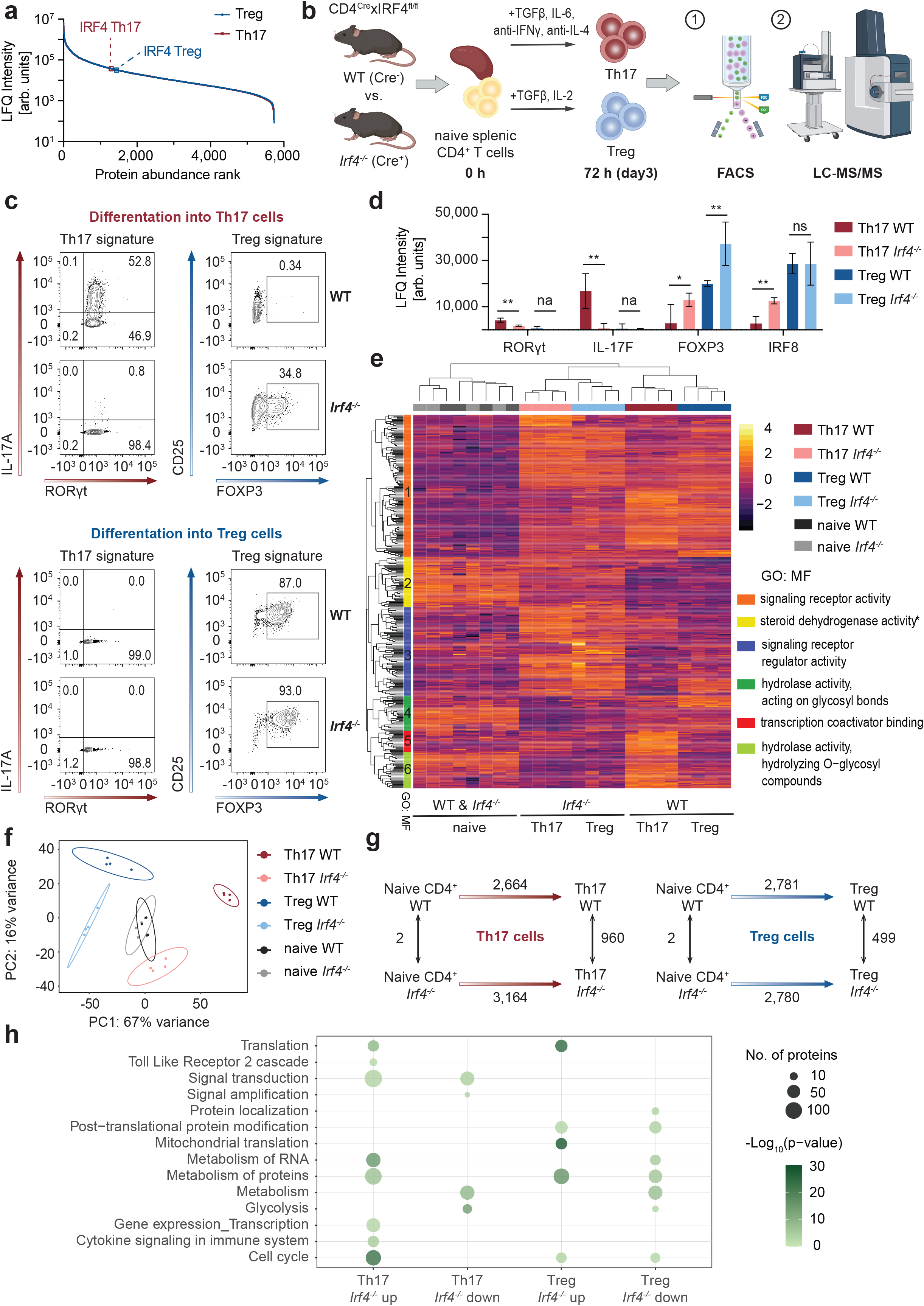
Proteomic analysis of Th17 and iTreg cells differentiated from naïve CD4^+^ T cells isolated from *Irf4^−/−^* and littermate control (WT) mice reveals marked changes in pathways required for cellular effector functions. (a) IRF4 expression levels in fully functional Th17 and iTreg cells from WT mice. (b) Schematic overview of the experimental setup for the analysis of *in vitro* polarized Th17 and iTreg cells. (c) FACS analysis of Th17 and iTreg cells grown in the presence of Th17/Treg-polarizing cytokines. Upon lack of IRF4, Th17 cells cannot develop into fully functional effector cells, showing impaired IL-17 expression while concomitantly displaying elevated FOXP3 levels. FACS analysis reveals similar (to slightly elevated) FOXP3 and CD25 levels in iTreg cells from *Irf4^−/−^* as compared to WT mice. (d) LFQ intensity values from LC-MS based proteome analysis for selected cell-type specific proteins. (e) Heatmap of proteins with significantly different expression levels in Th17 and iTreg cells from WT and *Irf4^−/−^* mice (*p-value* (limma) < 0.01, |log_2_(FC *Irf4^−/−^*/WT)| > 1). (f) PCA of protein abundances reveals a stronger separation of *Irf4^−/−^* Th17 cells from WT Th17 cells as for *Irf4^−/−^*iTreg cells from WT iTreg cells. (g) Numbers of differentially regulated proteins between naïve CD4^+^ T (0h) cells and fully differentiated Th17 and iTreg cells (72h) and *Irf4^−/−^* as well as WT mice. Numbers next to the arrows indicate the overall number of differentially expressed proteins (inclusive up- and downregulated proteins) between the different conditions (see also Supplementary Fig. 3). (h) Reactome pathway analysis of proteins that were upregulated upon differentiation (d3 as compared to d0) in an IRF4 dependent manner and/or in differentiated Th17 and iTreg cells from WT and *Irf4^−/−^* animals. Parts of panel (b) were created with Biorender.com and exported under a paid subscription.

In line with previous data^3^, proteomic analysis as well as flow cytometry-based analysis confirmed that IL-17 secreting Th17 cells cannot develop in the absence of IRF4 (Fig. 1c,d). Concomitant with strongly reduced IL-17 production, IRF4-deficient Th17 cells displayed significantly reduced expression of RORγt (Fig. 1d). In contrast, development of *in vitro* induced FOXP3^+^ (i)Treg cells from *Irf4^−/−^* naïve T cells was not affected. Interestingly, *Irf4^−/−^* iTreg cells showed significantly increased FOXP3 levels as compared to iTreg cells differentiated from WT mice confirming the repressive effect of IRF4 on FOXP3 expression (Fig. 1c,d). However, in-depth proteomic analysis revealed that *Irf4^−/−^* iTreg cells still differ markedly from WT iTreg cells regarding their protein expression pattern (see Fig. 1e,f,). Principal component analysis (PCA) of protein abundances demonstrated substantial differences between IRF4-deficient and IRF4-competent cells (Fig. 1f). While principal component 1 (PC1) reflects major changes between fully differentiated Th17 and iTreg cells, PC2 additionally separated *Irf4^−/−^*cells from their WT counterpart indicating a stronger divergence for *Irf4^−/−^* Th17 cells as compared to *Irf4^−/−^*iTreg cells. This is also reflected in the overall number of differentially expressed proteins: Upon differentiation from naïve CD4^+^ T cells (0 h) to fully functional Th17 and iTreg cells (72 h), a total of 2,664 proteins and 2,781 proteins changed their expression patterns, respectively (Fig. 1g, Supplementary Table 1). This number was also similar for iTregs from *Irf4^−/−^*mice. However, in *Irf4^−/−^*Th17 cells the number of proteins with differential expression levels upon differentiation (0 h vs 72 h) is markedly higher (3,164 proteins). This is also reflected in fully differentiated Th17 cells where around 960 proteins are differentially expressed between *Irf4^−/−^* Th17 cells and Th17 cells from WT animals. In iTreg cells, 499 proteins are differentially regulated between WT and *Irf4^−/−^* iTreg cells underlining the more prevalent role of IRF4 for Th17 cell development and functionality. *Irf4^−/−^*cells from both cell types show similar protein expression patterns (Fig 1e). Among the proteins that differ most in their expression level as compared to WT Th17 and iTreg cells (*p-value* < 0.01, log_2_(FC *Irf4^−/−^*/WT) > 1 or < −1), we found a strong enrichment of proteins associated with signal receptor (regulator) activity, transcription coactivator binding and hydrolase activity. Additionally, almost all the proteins are involved in either Th17 and/or iTreg cell differentiation displaying differential expression levels at 0 h and 72 h in the cells (Fig 1e). For example, (phospho)proteins associated with signaling receptor activity (cluster 1, Fig. 1e, Supplementary Fig. 1) are involved in protein (kinase) binding, the regulation of cell proliferation, motility and locomotion, and have been reported to be linked with (ab)normal adaptive (cell-mediated) immunity. This group includes IRF4 itself, cell surface receptors CD28, FAS, and CD69, adapter molecules (e.g., TRAF1), the TF NFAT5, MO4L2, RGCC, as well as kinases such as CDK6 and FADK1 (*Ptk2*). Proteins that are involved in signaling receptor regulator activity (cluster 3, Fig. 1e, Supplementary Fig. 2) are associated with the positive regulation of CD4^+^, alpha-beta T cell activation, regulation of metabolic processes, interspecies interaction between organisms including proteins involved in the defense against intracellular pathogens as well as (ab)normal T cell physiology and number. Examples are multiple TFs (including FOXP3, FOXP1, IRF1, IRF8, JunD, TCF-12), cell surface receptors LRB4A (*Lilrb4a*), CD166, and IL2RG which is involved in iTreg cell differentiation as well as cytokines involved in Treg cell development (e.g., XCL1), Nedd4 and other E3 ubiquitin-protein ligases.

We additionally performed Gene Ontology (GO) enrichment analyses exclusively for those proteins that are up-regulated during differentiation and displayed different expression levels between *Irf4^−/−^* and WT cells (Fig. 1h, Supplementary Fig. 3 and Supplementary Table 2). Upon lack of IRF4, proteins associated with protein binding, cellular metabolism, in particular protein and RNA metabolism, and pathways such as glycolysis were affected in both cell types, i.e., Th17 and iTreg cells. Previous studies have shown that metabolic pathways are ‘rewired’ upon activation and differentiation of resting lymphocytes shifting from oxidative phosphorylation to aerobic glycolysis fulfilling the bioenergetic demands during proliferation, differentiation and the biosynthesis of precursors^19^. IRF4-driven metabolic switch towards glycolysis has been demonstrated before for CD8^+^ T^20^ and Treg cells^9^, which is in line with our data. Distinct metabolic programs have also been associated with the polarization along the Th17 cell and Treg cell axis and oxidative phosphorylation has been correlated with a Treg cell phenotype^21^. Indeed, proteins involved in oxidative phosphorylation are up-regulated in *Irf4^−/−^*Th17 cells and down-regulated in *Irf4^−/−^* Treg cells (Supplementary Fig. 4). In addition, we also observed other marked differences between the two cell types that indicate a T cell subtype-specific role of IRF4 in the Th17 cell and iTreg cell cellular context. Namely, in iTreg cells, we found that mainly pathways regulating processes on “protein level” were affected such as, protein post-translational modification, protein localization and (mitochondrial) translation. In contrast, pathways and cellular processes that regulate (downstream) gene expression and transcription were impaired in *Irf4^−/−^* Th17 cells. Additionally, we found that the expression of proteins associated with signal transduction and immune cell functions (e.g., cytokine signaling, TLR2 cascade) were majorly affected by the lack of IRF4 in Th17 cells.

Interestingly, a subset of *Irf4^−/−^* CD4^+^ T cells cultured under Th17-inducing conditions displayed an increased expression of FOXP3 (34.8%) (Fig. 1c). Beside significantly increased FOXP3 levels, proteomic data of *Irf4^−/−^* Th17 cells revealed an elevated expression of other proteins associated with a more “Treg cell-like” phenotype (Fig. 1d), such as IRF8^22^, FOXP1^23^, CD166 (*Alcam*)^24^ and SMYD^25^ indicating a crucial role of IRF4 in Th17-Treg cell plasticity.

Our data further consolidate, in line with aforementioned studies, the importance and necessity of IRF4 for the development of fully functional Th17 and iTreg cells. However, the development and plasticity of immune cells depend on multiple factors and can be attributed to an interplay of different proteins, which directly or indirectly, as part of a large protein complex, contribute to cell fate.

### IRF4-mediated gene transcription is regulated by a subset-specific protein complex conserved across the Th17 cell and Treg cell axis

In an effort to discern which proteins act in concert with IRF4 and drive either a Th17 cell or Treg cell phenotype, we determined the cell-type specific IRF4 interactome in Th17 and iTreg cells by AP-MS (Fig 2a). In order to isolate IRF4 along with its protein interaction partners, we generated *in vivo* biotin-tagged IRF4 using the transgenic ROSA26^BirA^ mouse strain^16^ ubiquitously expressing the bacterial BirA protein biotin ligase from the ROSA26 locus. To generate *in vivo* biotin-tagged IRF4, we crossed ROSA26^BirA^ mice with transgenic mice expressing Avi-tagged IRF4 under the control of the endogenous *Irf4* promoter (IRF4^Avi-tag^ mice, Fig. 2b, Supplementary Fig. 5). The offspring (IRF4^Bio^ mouse) expresses biotinylated IRF4 at endogenous levels (Fig. 2c, Supplementary Fig. 6). IRF4^Bio^ mice are viable and fertile and show no overt phenotype. No difference could be observed between Th17 or Treg cells derived from naïve CD4^+^ T cells from IRF4^Bio^ or littermate control mice (at 72 h) indicating that the Avi-tag does not affect IRF4 activity and the expression of biotinylated IRF4 is not deleterious to Th17 or Treg cell development and function (Fig. 2d,e, Supplementary Figs. 8 and 9). As IRF4 is mainly located in the nucleus in fully differentiated cells (demonstrated exemplarily for Th17 cells in Fig. 2d, Supplementary Fig. 7), interactome analysis was conducted with the nuclear fraction. To efficiently reduce unspecific background, while at the same time preserving weak interactions, we applied a cell-permeable cross-linker, Dithiobis[succinimidyl propionate] (DSP) (Supplementary Fig. 10). Using DSP, protein-protein interactions can be preserved *in situ* prior to cell lysis and nuclear extraction (while cells are still alive)^26^. Moreover, we could markedly reduce unspecific binding as we were able to use more stringent wash conditions (Supplementary Fig. 10). Applying optimized digestion, wash and cross-linking conditions resulted in a reduced unspecific background as well as markedly diminished numbers of streptavidin peptides, while concomitantly detecting higher numbers of interaction partners including previously described IRF4 binding partners such as the TFs Ikaros and Aiolos^27^ at subcellular resolution (Fig. 2f, Supplementary Figs. 10 and 11, Supplementary Table 3).

**Fig. 2.**
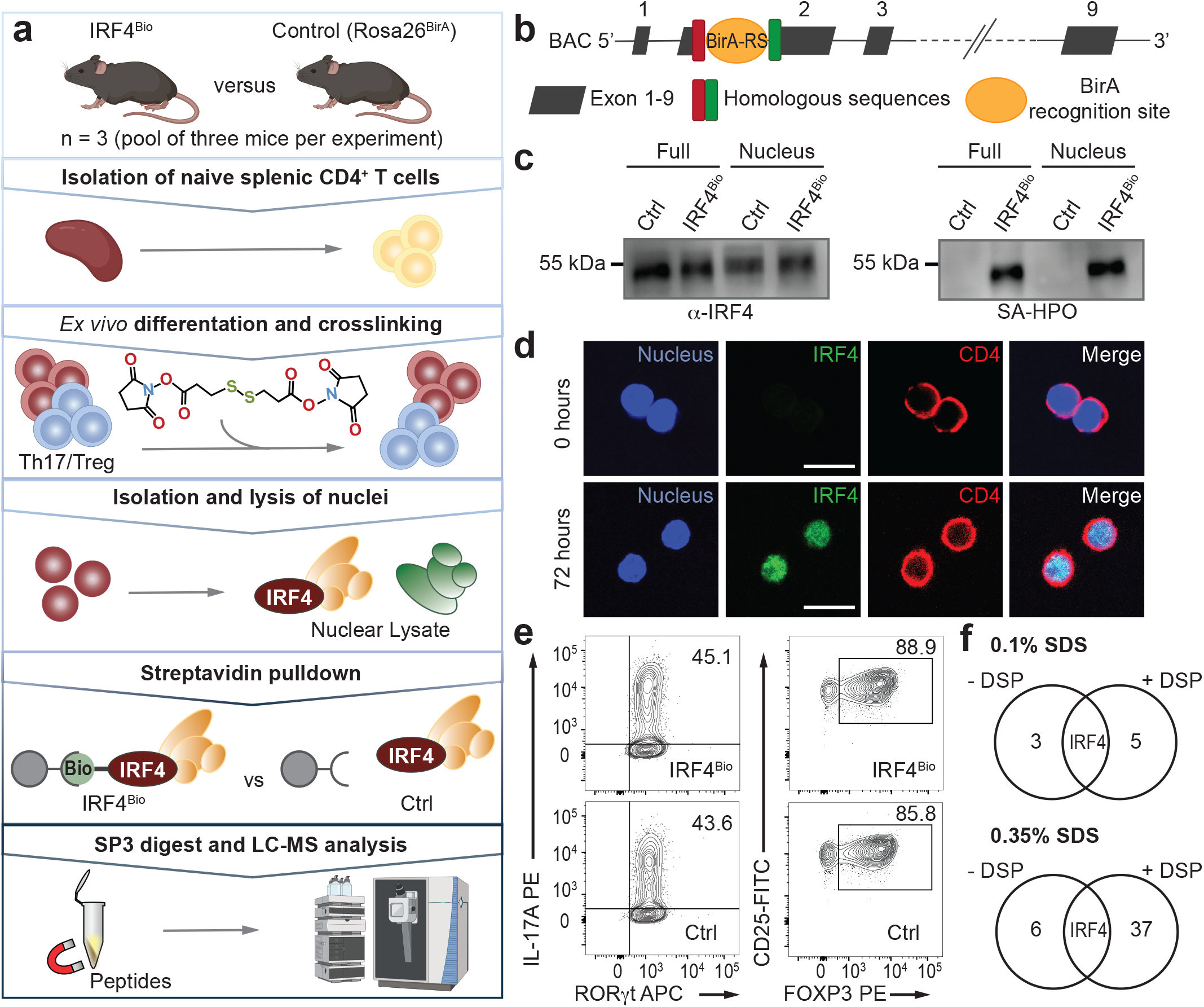
*In vivo* biotinylation and optimized affinity-purification of endogenous IRF4 complexes. (a) Graphical illustration of the optimized protocol for the isolation and MS-based characterization of IRF4 complexes derived from *ex vivo* differentiated Th17 and iTreg cells. (b) BAC transgene for the generation of the IRF4^Bio^ mouse. (c) Western blot analyses of full cell lysates and nuclear extracts demonstrate successful *in vivo* biotinylation of IRF4 (left panel: anti-IRF4 antibody; right panel: horseradish peroxidase-conjugated streptavidin (SA-HPO)) as exemplarily shown for Th17 cells. (d) Th17 cells show strong nuclear expression of IRF4 three days after isolation and *ex vivo* differentiation (white scale bar: 10 µm). (e) FACS analysis of murine Th17 and iTreg cells. (f) Chemical cross-linking combined with stringent washing conditions improves the quality and specificity of IRF4 pulldown experiments. Pulldown experiments were conducted with splenocytes isolated from IRF4^Bio^ and control mice. Parts of panel (a) were created with Biorender.com and exported under a paid subscription.

The optimized protocol was used to define the nuclear IRF4 protein interaction network in *ex vivo* generated Th17 and iTreg cells at day 3. Combining data from three independent AP experiments, we identified in total 440 IRF4 interactors across both cell types (Fig. 3, Supplementary Fig. 13, Supplementary Table 4). Functional network analysis of these IRF4 interactors drawing upon the STRING database of protein-protein interactions indicated a strong network enrichment, where 422 out of the 440 candidates (including IRF4) were assigned to a single interconnected network (Supplementary Fig. 13). Several proteins detected in the present dataset have been previously described to interact with IRF4 either in human and/or mouse models on the single molecule level such as Aiolos (encoded by *Ikzf3*), Ikaros (*Ikzf1*), STAT1, STAT3, VPS35, EP300, JunB and SATB1, as well as members of the Mi-2/nucleosome remodeling and deacetylase (NuRD) complex^27,28^ (Supplementary Table 4).

**Fig. 3.**
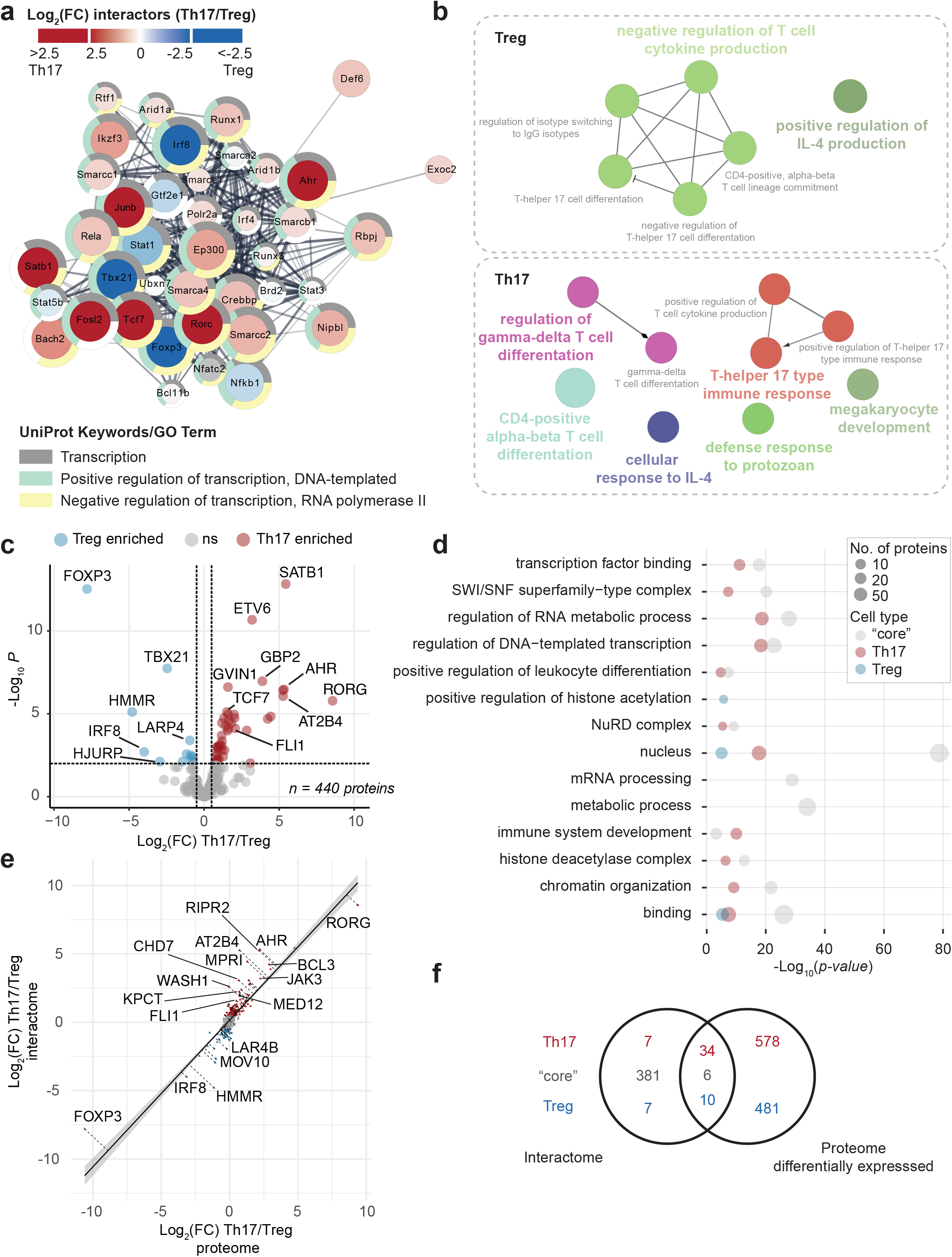
Characterization of the IRF4 interactome reveals novel players in IRF4-mediated gene transcription in fully functional Th17 and iTreg cells. IRF4 interactome analyses were performed using *ex vivo* generated Th17 and iTreg cells (72h). Naïve splenic CD4^+^ T cells from three mice were pooled per experiment prior to *ex vivo* differentiation. A total of three individual AP-LC-MS experiments per cell type was conducted. (a) STRING-based IRF4 interaction network. Displayed are interactors that have a direct functional and/or physical association with IRF4 (STRING DB default settings). The full IRF4 network is shown in Supplementary Fig. 13. Color scheme indicates enrichment either in Th17 (red) or iTreg cells (blue). (b) ClueGO analysis of IRF4 interactors that were significantly enriched either in Th17 or iTreg cells (*adj. p-value* < 0.01, |log_2_(FC Th17/iTreg)| > 0.5) (c) Volcano plot displaying differentially enriched IRF4 interactors in Th17 and iTreg cells (*adj. p-value* < 0.01, |log_2_(FC Th17/iTreg)| > 0.5). In total, 440 interactors were identified in the IRF4 complex across both cell types. (d) GO and reactome pathway analysis of IRF4 interactors. Grey: “core” IRF4 interactome, red: Th17, log_2_(FC Th17/iTreg) > 0.5, blue: iTreg log_2_(FC Th17/iTreg) < −0.5). (e) Comparison of log-transformed protein fold-changes at the interactome and at the proteome level between Th17 and iTreg cells. (f) Overlap of the IRF4 interactome with proteins that are differentially expressed in Th17 versus iTreg cells (full cellular proteome).

STRING network analysis indicated 39 direct associations, either functional and/or physical, between IRF4 and the proteins detected by AP-MS, including the Treg and Th17 lineage-defining TFs FOXP3 and RORγt, respectively (Fig. 3a,b). In total, we identified 53 proteins that were differentially enriched or exclusively detected in either the Th17 or iTreg cell IRF4 interactome (see Fig. 3c, 41 proteins enriched in Th17 cells, 12 proteins enriched in Treg cells). Beside the lineage-defining TFs, these include other characteristic (transcription) factors that are known to drive cell-type specific functions (Fig 3b). In iTreg cells, FOXP3, TBX21 and LAT1 (*Slc7a5*) negatively regulate Th17 cell differentiation, while promoting IL-4 production which can support suppressing immune responses by Treg cells^29,30^. Additionally, we found a strong enrichment of IRF8, a silencer of Th17 cell differentiation programs^31^, and of the hyaluronan-mediated motility receptor (HMMR) in the iTreg-specific IRF4 interactome. In Th17 cells, ARID5A, MALT1, RORγt, BCL-3, SATB1 promote CD4^+^ alpha-beta T cell differentiation and induce a Th17 cell-specific immune response (Fig. 3b, Supplementary Table 5). SATB1, for example, has been described to differentially regulate gene expression in Th17 cells promoting the pathogenic effector program of encephalitogenic Th17 cells^32^. The NF-κB regulator MALT1, which to our knowledge has not yet been described as part of the IRF4-regulated transcription network determines the encephalitogenic potential of inflammatory Th17 cells in experimental autoimmune encephalomyelitis, a mouse model of multiple sclerosis^33^. Other IRF4 interacting proteins enriched in Th17 cells, steering lineage-specific gene expression, include CBP/EP300, TCF7, AHR, Fosl2, CHD7, ZN609, BACH2, JunB, and ETV6^7,34^. Whereas JunB has been described to co-localize with IRF4 at the *Il17* (and *Il21*) promoter^35^ and is essential for IL-23-dependent pathogenicity of Th17 cells^36^, other factors such as TCF7, FLI1, BACH2, GTF2I have not yet been associated with the IRF4 transcriptional regulatory network in Th17 cells. TCF7 (transcription factor 7 or T cell factor 1, TCF1), a major positive regulator of chromatin accessibility required for early T cell development, has been linked with a stem-like Th17 cell state^37^ and Th17 longevity^38^. Interestingly, TCF7 expression is reduced and fine-tuned by FOXP3 to drive Treg cell-specific repression of chromatin accessibility and gene expression^39^. Although BACH2 has not yet been described as a direct interaction partner of IRF4 in fully differentiated Th17 cells, it has been shown that BACH2 directly steers and attenuates TCR-induced IRF4-dependent transcription in Treg cells^40^.

Interestingly, a large number of proteins was identified in the IRF4 interactome of both cell types indicating common regulatory mechanisms in IRF4-steered gene regulation across the two functionally opposing T cell subsets (Fig 3c). For proteins identified in the IRF4 “core” interactome in both cell types, i.e. that display no significant enrichment in either iTreg or Th17 cells, GO analysis revealed a strong enrichment of nuclear proteins involved in DNA-templated regulation of transcription, many of them displaying either TF, DNA or poly(A) RNA binding capabilities (Fig. 3d, Supplementary Fig. 14, Supplementary Table 5). Moreover, proteins involved in chromatin organization as well as proteins associated with (m)RNA processing and regulation of (RNA) metabolic processes represent the “core” IRF4 interactome. The majority of proteins identified in the IRF4 “core” interactome also displayed no differential expression between the two cell types, i.e. in Th17 and iTreg cells, on full proteome level (Fig. 3e,f, Supplementary Table 6). Among respective proteins, we also identified several members of the ATP-dependent chromatin remodeling complex NuRD, which operates as a global modulator of transcription acting broadly at many enhancers and promoters to dampen and fine-tune active gene expression^41^. By targeting the NuRD complex, (IRF4-)Ikaros and Aiolos influence its genomic localization and functional activity^27,42^. Additionally, we detected the chromatin-remodeling SWI-SNF (switch/sucrose nonfermenting) complex. As transcriptional activator, SWI-SNF has been reported to have an antagonistic role to NuRD on common regulatory elements^41^. Both complexes are required during cellular commitment ‘decisions’ regulating both B and T cell differentiation (and together with Ikaros form the so-called PYR-complex). Data derived from T cells suggest that NuRD/Mi-2 can determine the transcriptional activity of TFs in a stage-specific manner and that concomitant interactions between functionally opposing chromatin-regulating machineries are an important mechanism of gene regulation during lineage determination^43,44^. Other “core” interactors are protein kinases, such as CDK7, CDK2, CDK1 and CDK9 involved in cell cycle control as well as RNA polymerase II-mediated RNA transcription, regulation of gene expression and DNA-templated transcription. CDK9, for example, steers cytokine inducible transcription networks (such as IL-6-inducible STAT3 signaling) whereas CDK2, by phosphorylating FOXP3, negatively regulates its transcriptional activity and protein stability^45^.

### ChIP-Seq analysis reveals a strong enrichment of motifs from IRF4 binding partners

To identify target genes regulated by the IRF4 complexes in Th17 and iTreg cells, we additionally performed streptavidin-mediated chromatin precipitation coupled with deep sequencing (Bio-ChIP-Seq) (Fig. 4, Supplementary Table 7). ChIP-Seq analysis of IRF4 in Th17 cells revealed 21,884 IRF4 binding regions annotated to 10,918 unique genes. Most peaks in the Th17 cells (41.9 %) were found in the promoter region (up to 3 kb downstream of the transcription start site (TSS)), followed by intronic binding (26.2 %) and distal intergenic binding (26.0 %) (Fig. 4b). In iTreg cells, a total of 10,189 binding regions were detected corresponding to 5,789 genes. Binding of IRF4 was most prevalent in introns (32.9 %) followed by promoter binding (31.9 %) and distal intergenic binding (31.0 %) (Fig. 4b). A total of 5,314 genes showed IRF4 binding in Th17 and iTreg cells, while 5,604 genes were exclusively bound in Th17 cells and 475 genes in iTreg cells. The high number of genes regulated by the IRF4 complex in Th17 cells along with the large number of associated promoter regions is in line with previous findings and further underlines the more prevalent role of IRF4 in Th17 as compared to iTreg cells. Moreover, in line with previous studies^8,46^, we could confirm that IRF4 directly binds within the promoter regions of the *Il17a* and *Rorc* genes. Of note, distinct IRF4 binding sites were detected within in the distal intergenic, intron and the promoter regions of *Il17a* and *Rorc* exclusively in Th17 cells (Fig. 4a). In both cell types, we could not detect any binding within the *Foxp3* locus.

**Fig. 4.**
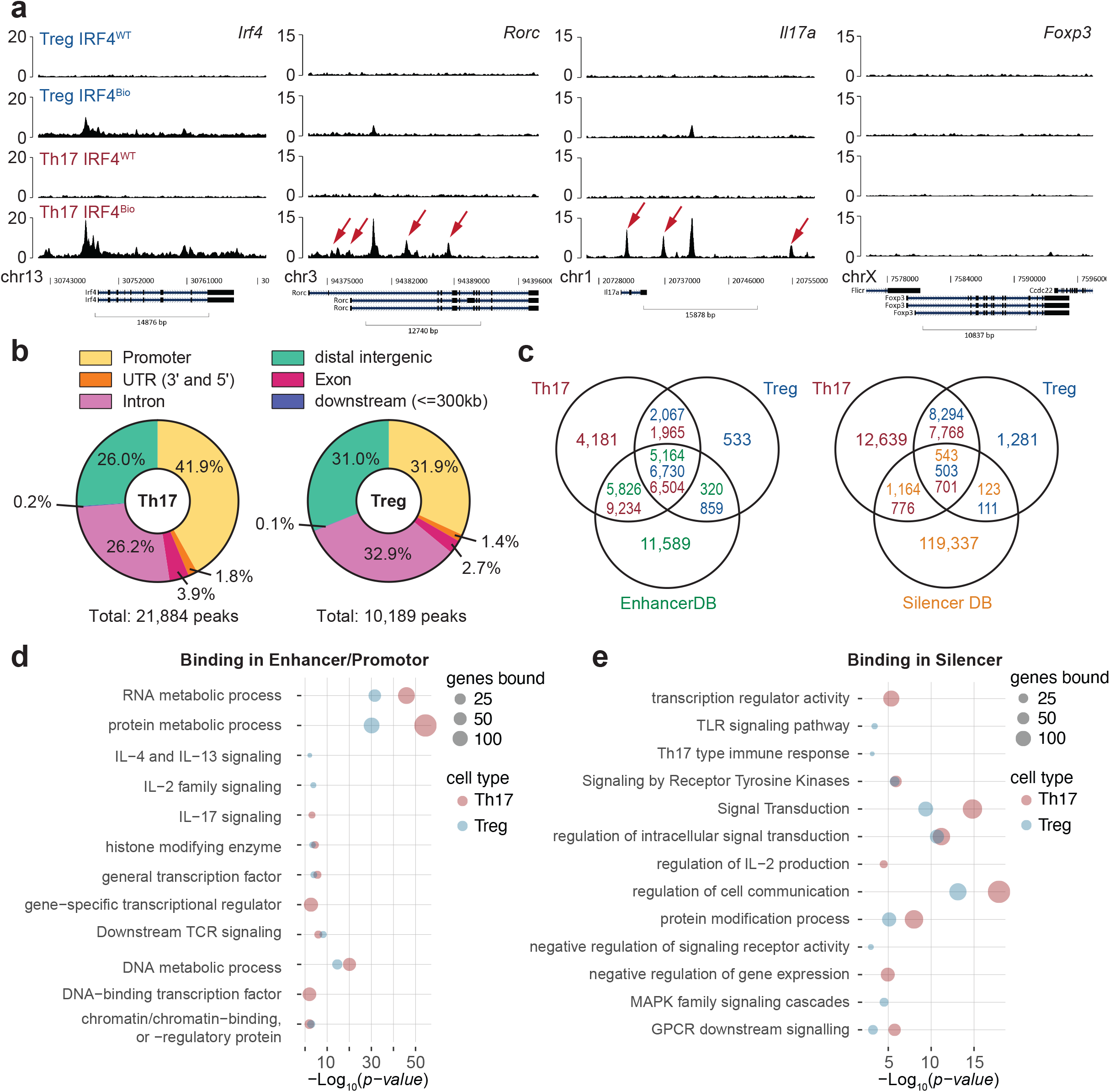
IRF4-ChIP-Seq analysis reveals twice the number of regulated genes in Th17 as compared to iTreg cells. Bio-ChIP-seq analysis of IRF4 binding was performed in Th17 and iTreg cells differentiated *ex vivo* from naïve splenic CD4^+^ T cells of IRF4^Bio^ mice. Cells derived from ROSA26^BirA^ (marked as IRF4^WT^) served as control. (a) ChIP-seq binding tracks for selected loci in Treg and Th17 cells at 72 h. (b) Distribution of identified binding sites in Th17 and iTreg cells. (c) Overlap of identified binding sites with EnhancerDB^47^ and SilencerDB^48^ knowledge bases. (d,e) GO and Reactome pathway analysis of genes bound by IRF4 in (d) enhancer/promoter and (e) silencer regions.

Approximately, 72% of the Th17 peaks mapped to predicted enhancer structures (10,990 unique enhancers) of the CD4^+^ cell data in the Enhancer Atlas2.0^47^, while 1,477 peaks (6.7 %) were associated with gene expression silencing (corresponding to 1,707 silencer regions, SilencerDB^48^). In iTreg cells, 7,589 peaks (74.5 %) showed an overlap with the enhancer database (mapping to 5,484 enhancer structures), among these 88.7 % were shared with Th17 cells. Out of the 614 peaks (6.0 %) mapping to unique silencer structures in iTreg cells, 543 (88.4 %) were also detected in Th17 cells. Applying functional enrichment analysis, we investigated which genes are regulated by the IRF4 complex in Th17 and iTreg cells. We found that expression of genes associated with RNA, DNA and protein metabolic processes are promoted by IRF4 in both cell types (Fig 4d, Supplementary Table 8). This ties in nicely with the proteomic data, where we observed a de-regulation of RNA, DNA and protein metabolic pathways upon lack of IRF4. In addition, genes promoting the expression of general TFs, histone modifying enzymes and chromatin/chromatin-binding, or -regulatory proteins were found to be positively regulated by IRF4 in both T cell types. Elevated gene expression for gene-specific transcriptional regulator or DNA-binding TFs was only seen in Th17 cells. This is also in line with the proteomic dataset where we mainly observed an impaired expression of proteins associated with transcription in Th17 but not iTreg cells (Fig. 1). The expression of genes associated with downstream signaling of the TCR is enhanced by IRF4 in both cell types. However, genes related to IL-17 signaling are only promoted in Th17 cells by IRF4, while in iTreg cells IL-2 family signaling as well as IL-4 and IL-13 signaling were increased (Fig 4d). Genes that are silenced by IRF4 in both cell types were mainly associated with signal transduction and the regulation of cell communication. Interestingly, in iTreg cells IRF4 complexes down-regulate Th17 type immune responses as well as TLR and MAPK signaling, while in Th17 cells genes related with the regulation of IL-2 production, transcription regulator activity and the negative regulation of gene expression were silenced (Fig. 4e, Supplementary Table 8). Despite the large number of commonly targeted genes across Th17 and iTreg cells, these data further demonstrate that IRF4 additionally acts in a context-dependent and hence cell-type specific manner.

To identify TF binding sites within the immunoprecipitated DNA fragments, we conducted a motif analysis using the MATCH algorithm^49^ and the murine TRANScription FACtor (TRANSFAC) database^50^ (Fig. 5, Supplementary Table 7). This revealed binding motifs from 133 unique proteins with the majority, i.e. 129 associated proteins, reported for both cell types (Fig. 5a,b). Interestingly, the binding motif of AHR was exclusively identified in Th17 cells, while the FOXP3, IRF8 and NR2C2 motifs were detected only in iTreg cells. The best matching count matrix for IRF4 was described in 21,846 peaks (99.83 %) of the Th17 ChIP and in 10,048 peaks (98.62 %) of iTreg cells confirming efficient isolation of IRF4 DNA binding regions. In addition, we applied the MEME-ChIP algorithms^51,52^ to search for enriched motifs. In total, 42 motifs were identified to be enriched in the DNA fragments of Th17 cells and 21 motifs in iTreg cells. To identify TFs associated with the enriched motifs, we used the Tomtom algorithm^53^ for motif mapping against the murine TRANSFAC database revealing 171 matching proteins (for 33 motifs) in Th17 cells and 206 unique proteins (for 17 motifs) in iTreg cells (Fig. 5a,b). Eight motifs in the Th17 ChIP and four motifs from the iTreg ChIP were identified as enriched but could not be mapped to any TF from the TRANSFAC database (Fig. 5c). Enriched motifs mapped to 152 proteins that were reported for both cell types. Out of these 42 TFs were also identified by the MATCH/TRANSFAC algorithm in both iTreg and Th17 cells, including IRF4, FLI1, STAT3, BACH2, STAT1, and JunB. All those proteins were also detected in the IRF4 interactome in both cell types indicating that they act and transcribe genes in concert with IRF4 in fully differentiated iTreg and Th17 cells. Although the motif of the IRF4 interactor EP300 was identified by MATCH/TRANSFAC in both cell types, enrichment was only in reported in Th17 cells. Motifs of IRF8 and NR2C2 were identified as enriched in both cell types but were only found in iTreg cells by the MATCH/TRANSFAC algorithm. Additional motifs identified by MATCH/TRANSFAC mapped to SATB1, HDAC1, RFX1, NFKB1, CBP, GTF2I and TCF7, which were also all identified in the IRF4 interactome.

**Fig. 5.**
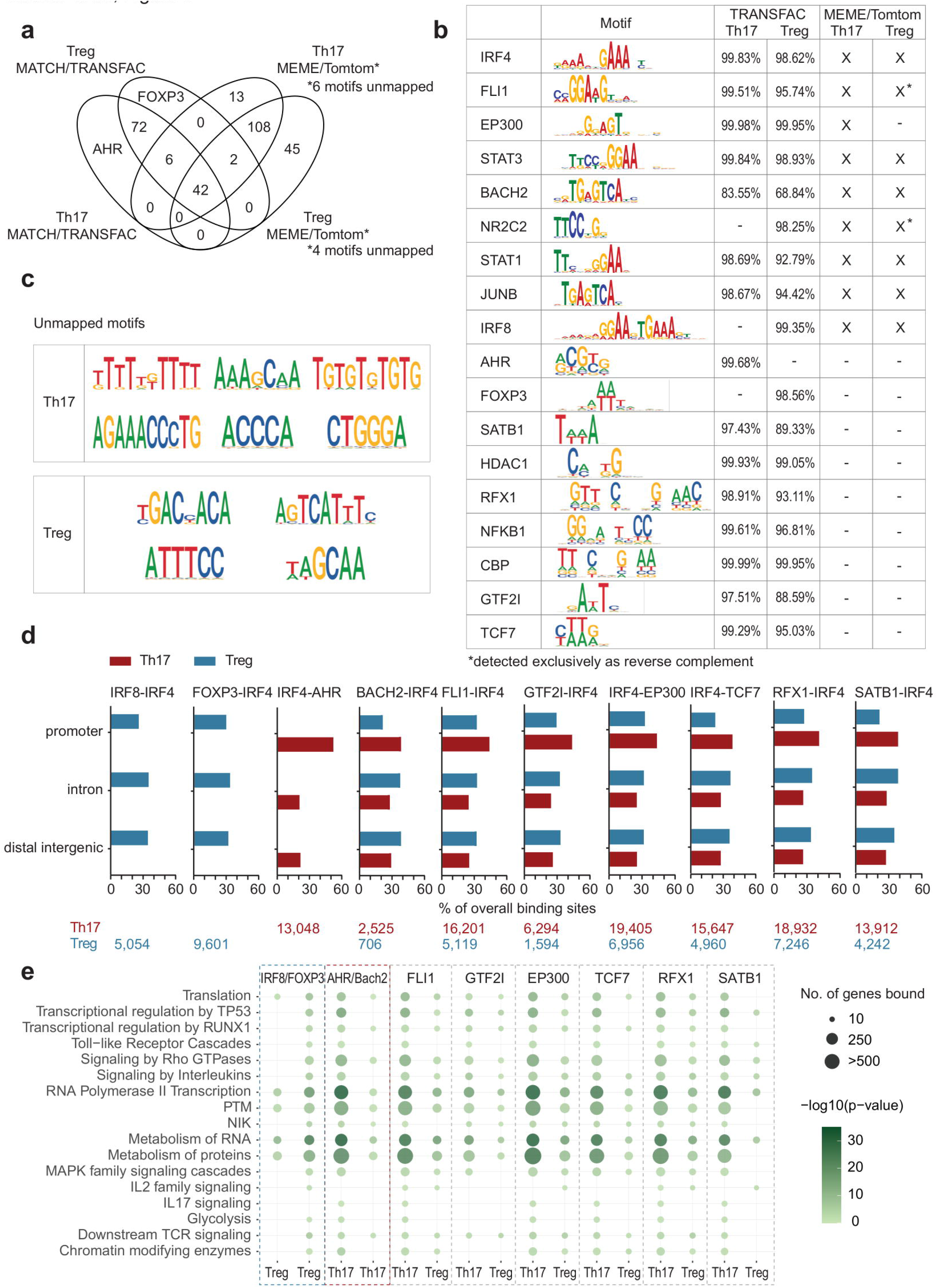
Motif analysis reveals binding sites of IRF4 interaction partners in genes encoding molecules involved in regulating effector and metabolic functions of Th17 and iTreg cells. Bio-ChIP-seq analysis of IRF4 binding was performed in Th17 and iTreg cells differentiated *ex vivo* from naïve splenic CD4^+^ T cells of IRF4^Bio^ mice. (a) Overlap of motifs identified by MATCH/TRANSFAC and MEME/Tomtom in Th17 and iTreg cells. (b) Binding motifs of IRF4-interactors were identified and enriched in the IRF4-ChIP-Seq data. (c) Novel unmapped motifs identified in Th17 and iTreg cells. (d) Distribution of combined binding motives across the peaks identified by IRF4-ChIP-Seq (IRF4 and indicated interactor within a proximity of 5 bp). Number of peaks where combined motifs were detected is indicated below the bar plots. (e) Reactome analysis of genes regulated by IRF4 and its interactors in Th17 and iTreg cells.

IRF4, which on its own displays only weak DNA-binding affinity, typically associates with other TFs, e.g., PU.1 and Spi-B in B cells or AP1 in CD4^+^ T cells, resulting in the recruitment of the (heterodimeric) complex to composite motifs (ETS–IRF or AP1–IRF4 composite elements, referred to as EICE/AICE motifs) in the genome^8,54,55^. To further investigate how IRF4 and its partners regulate gene transcription, we also conducted a combined motif analysis to identify motifs in close proximity (up to 5 bp distance) to each other (Fig. 5d, Supplementary Table 7) and found that nearly all previously detected motifs of IRF4 interactors were located in the immediate vicinity of the IRF4 motif. Additionally, the motifs of IRF4 interactors were also detected in close proximity to each other (Supplementary Table 7). In line with the single motif analysis, FOXP3-IRF4 and IRF8-IRF4 composite motifs were exclusively identified in the iTreg ChIP-Seq dataset, while IRF4-AHR composite elements were exclusively reported for Th17 cells. The FOXP3-IRF4 motif is the most prominent composite motif in iTreg cells (around 9,601 associated peaks), followed by RFX1-IRF4 and IRF4-EP300 motifs. In Th17 cells, we found multiple composite motifs that were associated with a high number of target genes, including IRF4-AHR, FLI-IRF4, IRF4-EP300, IRF4-TCF7, RFX1-IRF4 and SATB1-IRF4 (13,000 to >19,000 peaks). In addition, most of these coincident binding peaks are located in promotor regions suggesting a driving role for respective IRF4 interactors in maintaining Th17 functionality. This could be further confirmed by GO analysis. Target genes containing composite motifs and associated with IL-17 signaling and glycolysis are co-regulated by IRF4 and AHR, FLI1, EP300, TCF7, RFX1 and SATB1 in Th17 cells (Fig. 5e, Supplementary Table 10). Additionally, we found that genes associated with TLR signaling are mainly targeted in Th17 but not in iTreg cells. In general, GO analysis revealed similar gene regulatory patterns for IRF4 complexes between iTreg and Th17 cells. However, it should be noted that the proportion of peaks positioned in intronic regions was much higher in iTreg cells as compared to Th17 cells, where IRF4 complexes and heterodimers favor promoter binding.

To investigate which proteins are directly affected by the IRF4-driven gene regulation, we integrated the information derived from the interactome, ChIP-seq and *Irf4^−/−^* proteome analyses (Fig. 6). To this end, we analyzed which proteins correspond to genes that are targeted by IRF4 heterodimers (i.e., multimers) and are affected by the lack of IRF4 (Fig. 6a, Supplementary Figure 15, Supplementary Table 1). In total, we identified 1,017 proteins in Th17 cells that showed de-regulated expression patterns upon differentiation (d0 vs d3) and/or in fully differentiated Th17 cells (d3) between WT and *Irf4^−/−^* animals and were directly regulated on the gene level by IRF4 and distinct interactors (binding either in ‘promoter + enhancer’ or ‘promoter + silencer’ region). In iTreg cells, this was the case for 280 proteins (Fig. 6a, Supplementary Fig. 15). In both cell types, IRF4 co-operates with multiple TFs to regulate gene transcription resulting in different sets of expressed proteins. In iTreg cells IRF4 mainly acts in concert with FOXP3 and EP300, while in Th17 cells EP300, RFX1, FLI1 and AHR play a dominant role. Among the proteins regulated by the IRF4 complex, we detected a large number of TFs including IRF4 interactors, as well as a large number of proteins associated with cellular metabolism and immune response (Fig. 6c-e). Nine proteins that were targeted by IRF4 in both cell types displayed similar protein expression patterns upon differentiation and lack of IRF4 indicating common regulatory mechanisms. These include the histone deacetylase 7, which enhances the transcriptional repressor activity of FOXP3 (by similarity, UniProt), the GTPases RALB and GBP4, the mitochondrial NAD-dependent malic enzyme 2 (ME2) crucial for mitochondrial pyruvate and energy metabolism^56^, TAXB1 and LAPM5 involved in the (negative) regulation of T cell activation and NF-kappa-B signaling, as well as the RNA helicase DDX24 and the protein S10AA (Fig. 6d).

**Fig. 6.**
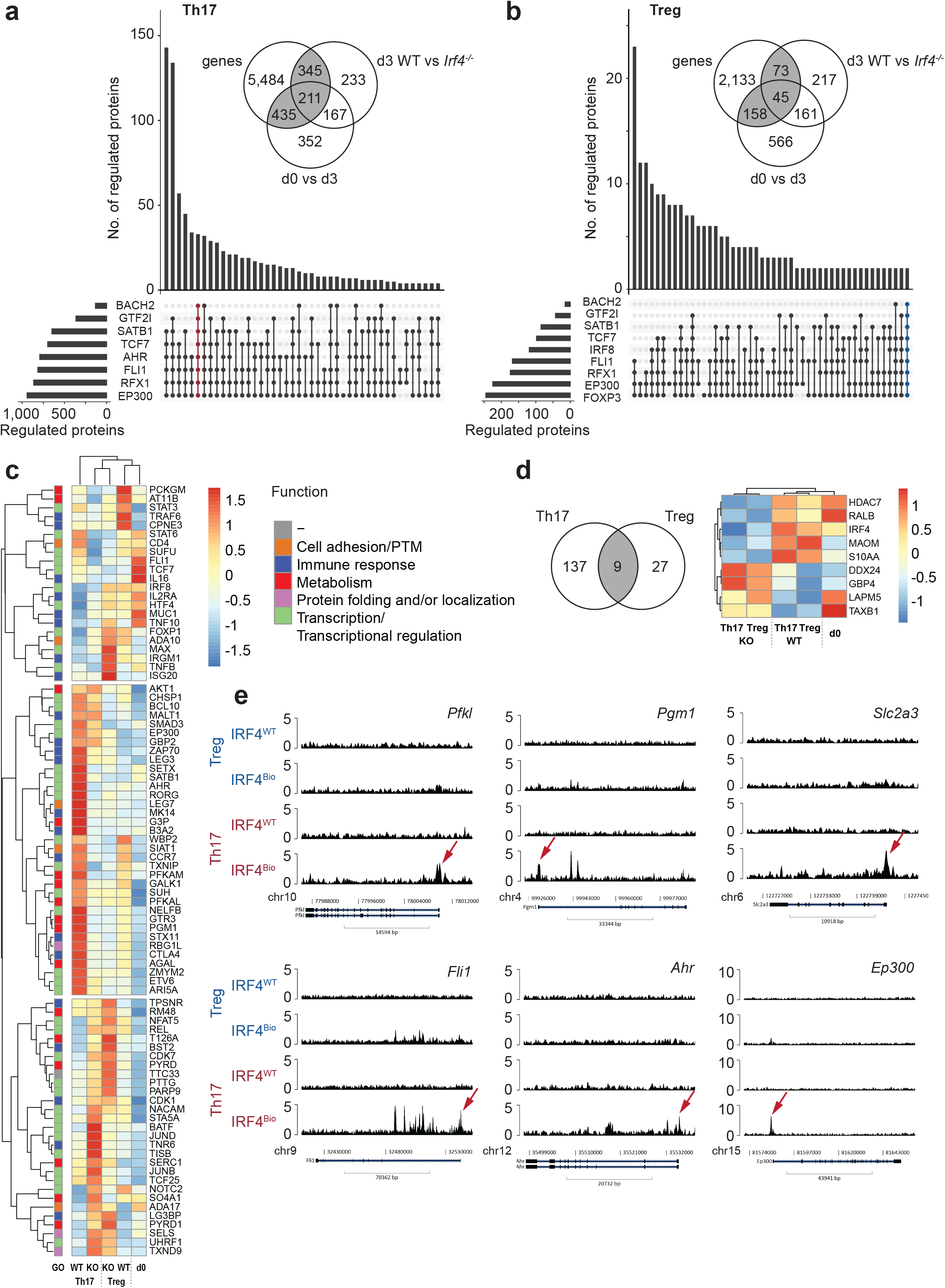
Genes regulated in fully differentiated Th17 and iTreg cells display altered proteomic patterns in *Irf4^−/−^* animals. Number of genes regulated by IRF4 and distinct interactors in (a) Th17 and (b) iTreg cells. Insets show an overlap of regulated genes (filtered for IRF4 binding in ‘promoter + enhancer’ or ‘promoter + silencer’), IRF4-dependent differentially expressed proteins at d0 and d3, as well as differentially expressed proteins in Th17 and iTreg cells of *Irf4^−/−^*and littermate control mice (see also Supplementary Fig. 15) (c) Protein expression patterns of selected candidates regulated by the IRF4-complex that display differential protein levels upon IRF4 knock-out. (d) Genes (proteins) regulated by IRF4 that show the same patterns in Th17 and iTreg cells. (e) ChIP-seq binding tracks for selected loci in iTreg and Th17 cells at 72 h. Red arrows indicate enhancer regions.

In Th17 cells we found a large number of proteins associated with transcriptional regulation and Th17 cell differentiation, as well as glucose metabolism (Fig. 6c,e, Supplementary Fig. 16). Interestingly, TFs and transcriptional regulators that are part of the IRF4 interactome, such as AHR, EP300, FLI1, ETV6, RORγt, TCF7, SATB1, MALT1, ZMYM2, ARI5A, and SUFU, are transcriptionally regulated by the IRF4 complex itself in Th17 cells and display low protein expression levels upon the lack of IRF4 indicating a positive feedback regulation for the respective gene products (Fig. 6c,e). In Treg cells, gene expression of proteins involved in protein localization, lipid and rRNA binding/translocation as well as transcriptional regulation is directly steered by the IRF4 complex. Examples of transcriptionally regulated proteins that display aberrant expression upon lack of IRF4 in iTreg cells are TRAF6 required for posttranslational FOXP3 regulation^57^, Copine-3 and the phospholipid-transporting ATPase IF, which play a role in cell migration and the maintenance of membrane lipid asymmetry in the endosomal compartment. Taken together, these data reveal distinct IRF4-driven transcriptional programs in functionally opposing Th17 and iTreg cells, where IRF4 in concert with specific interactors regulates the expression of genes crucial for full Th17 and iTreg cell functionality.

## DISCUSSION

IRF4 is crucial for the differentiation and fate determination of different CD4^+^ T cell subpopulations, including pro-inflammatory Th17 cells as well as the functionally opposing group of immunomodulatory Treg cells^3,10,58,59^. Over the past decade multiple studies have elucidated IRF4-mediated transcriptional programs in (differentiating) Th17 and Treg cells. However, to our knowledge, no study investigated IRF4 directly comparing fully differentiated Th17 and Treg cells providing an unbiased view on the molecular mechanisms of IRF4-steered gene expression and IRF4 interactions in respective cell types.

Integrating data derived from IRF4-ChIP-Seq, interactome and discovery-based proteome analyses from WT and *Irf4^−/−^* animals, we could, consistent with previous research^3^, demonstrate that IRF4 plays a more prevalent role in Th17 as compared to (i)Treg cells. While CD25^+^ FOXP3^+^ expressing immunomodulatory Treg cells could develop from *Irf4^−/−^* naïve CD4^+^ T cells, the development of Th17 cells was markedly impaired in the absence of IRF4. We could not detect any IL-17 secreting cells by FACS analysis and proteomic characterization of *Irf4^−/−^*cells resulted in a higher number of dysregulated proteins in Th17 as compared to iTreg cells. This was further affirmed by ChIP-seq analysis, which revealed twice as many IRF4 binding sites, especially in the promoter regions, in Th17 as compared to iTreg cells.

Integrating the information derived from proteome and ChIP-Seq analysis, we found some common IRF4-driven regulatory mechanisms as well as cell-type specific ones. Our data revealed, for example, that IRF4 regulates RNA and protein metabolic processes as well as pathways associated with cellular energy metabolism in both, Th17 and iTreg cells. This is in accord with previous studies where IRF4-mediated metabolic programming has been described for different effector T cell subsets, such as CD8^+^ T cells^20^, Th1^60^ and Treg cells^9^. Upon T cell activation glycolysis is upregulated in an IRF4-dependent manner^20,60,61^ and the expression of proteins that promote uptake of nutrients, e.g., glucose transporter GLUT3 (*Slc2a3*), also detected in the present dataset, are directly controlled by IRF4. Interestingly, proteins associated with oxidative phosphorylation, a process correlated with a Treg cell phenotype, were up-regulated in *Irf4^−/−^* Th17 cells and down-regulated in *Irf4^−/−^* Treg cells pointing to cell-type specific IRF4-dependent regulatory mechanisms. Other cell-type specific effector proteins detected in the present dataset and controlled by IRF4 are pro-inflammatory cytokines in Th17 cells. The role of IRF4 in pro-inflammatory cytokine production, especially IL-17 signaling, has been described previously^8,46^ and linked with Th17-mediated diseases, like EAE^3^ and colitis^62,63^.

Moreover, we detected a large number of genes and proteins associated with gene transcription and transcriptional regulation that were directly controlled by IRF4 in Th17 but not iTreg cells further underlining the cell-type specific regulatory functions of IRF4. In case of iTreg cells, our ChIP-seq and proteomic data revealed IRF4-dependent transcriptional regulation of genes associated with IL-2 and IL-4 signaling. When co-expressed, these two cytokines enhance Treg proliferation and immunosuppressive function, e.g., IL-10 secretion^64^. IRF4-dependent regulation of IL-2 and IL-4 production has been investigated previously in naïve and effector/memory CD4^+^ T cells from *Irf4^−/−^* mice demonstrating that IRF4-dependent IL-4 regulation takes place in a context-dependent manner^65^. Moreover, it has been previously shown that IRF4 maintains the expression of genes associated with cell cycle progression concomitantly repressing pro-apoptotic genes in multiple myeloma and loss of IRF4 is linked to decreased proliferation *in vitro* and reduced tumor growth *in vivo*^66–68^. In our dataset, we found that IRF4 is also involved in cell cycle regulation in iTreg as well as Th17 cells suggesting a similar regulatory role for these cell types.

Interestingly, *Irf4^−/−^* cells grown under Th17-skewing conditions expressed the Treg-specific TF FOXP3 along with other molecules that are important for Treg homeostasis and anti-inflammatory function, such as IRF8, FOXP1 and SMYD3 demonstrating the crucial role of IRF4 for Th17 development. FOXP1 stabilizes FOXP3 expression and is critical for Treg suppressive functions^23^, while the histone methyltransferase SMYD3 controls the stability of iTreg cells on an epigenetic level^25^. Loss of either molecule results in pro-inflammatory immune responses^23,25^, including the expression of Th17-associated genes, such as *Il17a*, *Il21* and *Rorc*, under Treg-skewing conditions^25^. Expression of the IRF4 homologue IRF8 is regulated by FOXP3 and it has been demonstrated that IRF8 suppresses Th17 cell development by inhibiting RORγt expression and subsequent IL-17 production^31^. Of note, we also identified IRF8, along with FOXP3, in the IRF4 interactome in iTreg cells further supporting the diametrically opposite effects of IRF4 and IRF8 as suggested previously^22^, with IRF8 promoting the expansion of immune-suppressive mechanisms in Treg cells^69^.

To further unravel the underlying cell type-specific IRF4-steered regulatory mechanisms, we performed an unbiased IRF4 interactome analysis directly comparing Th17 and iTreg cells using a novel mouse strain. We identified a large number of common IRF4 interactors (286 proteins associated with the IRF4 “core” interactome) in both cell types, including members of the NuRD and SWI/SNF superfamily-type complexes and other interactors associated with the regulation of DNA-templated transcription and mRNA processing while other proteins were subtype specific. An overlap with literature-curated IRF4 interacting proteins verified in total 41 proteins (Supplementary Table 3), but also revealed novel interactors, such as FLI1, BACH2, AHR, GTF2I, RFX1, and MALT1, among many others, which were not yet described. Moreover, the majority of known IRF4 interactors have not yet been reported in the Th17/Treg context, but have been associated with other (immune) cell types, e.g. macrophages^70^, plasma cells^71^ or other T cell subsets^14^. To our knowledge, in Treg cells only a FOXP3-IRF4 interaction was demonstrated, which regulates the expression of genes essential for the anti-inflammatory character of this cell type, e.g. *Icos*, *Il10* or *Grmb*^10,11^. An interaction of IRF4 with JunB^8,54^ or STAT3^3,11^ in Th17 cells was described by co-localization and overlapping DNA-binding regions of those TFs at Th17-critical genes, such as *Rorc*, *Il17a*, *Il17f* and *Il21*^46,72^, however, no physical interaction could be verified by reproducible immunoprecipitations^8,72^. Data from our unbiased interactome analysis matched with these results and revealed reproducibly IRF4-JunB and IRF4-STAT3 interactions in Th17 cells.

As a weak DNA-binder, IRF4 typically associates with cell-type specific partners to be recruited and bind distinct composite sequence elements^54,55^, such as the AICE and ISRE composite elements in T and B cells as well as the EICE and ZICE motifs in B cells. Conducting motif analysis, we detected multiple motifs in our dataset mapping to IRF4 interactors, e.g., for JunB, STAT3, EP300 FLI1, TCF7, AHR, IRF8, FOXP3, SATB1, etc. Interestingly, combined motif analysis revealed multiple composite motifs for IRF4 and its binding partners that have not been described before. While IRF4 combinatorial motifs, i.e., for BACH2, FLI1, GTF2I, EP300, TCF7, RFX1, and SATB1 were detected in both cell types, we exclusively identified “IRF8-IRF4” and “FOXP3-IRF4” composite motifs in Treg and “IRF4-AHR” elements in Th17 cells. Linking the genes where IRF4 complexes bind either in ‘promoter + enhancer’ or ‘promoter + silencer’ regions with the *Irf4^−/−^* proteome dataset, we could identify a set of proteins transcriptionally regulated by IRF4 and distinct interactors, which display aberrant expression also at the proteome level. Among these, we found proteins associated with immune response, metabolism as well as a large number of transcriptional regulators. Interestingly, among the transcriptional regulators, we identified several IRF4 binding partners, e.g. AHR, FLI1, EP300, TCF7, GTF2I, RFX1, SATB1, etc., that bind in concert with IRF4 in their own promoter region suggesting a positive feedback regulation for the respective molecules in Th17 cells. Most of these factors have been associated to play crucial roles in the Th17/Treg equilibrium. For example, AHR, exclusively detected as IRF4 interactor in Th17 cells, can induce anti-inflammatory Treg as well as pro-inflammatory, pathogenic Th17 cells. Activated AHR regulates gene expression either binding directly to a specific consensus motif (5L-TNGCGTG-3L) or by association with other TFs, e.g., RelA, STAT proteins or members of the SWI/SNF complex^73,74^, detected also in the present dataset. In Th17 cells, AHR expression is induced by STAT3 and strongly promotes the Th17 cell fate by boosting IL-17A, IL-17F and IL-22 production and by interfering with Treg cell pathways limiting, for example, STAT5 and STAT1 signaling or inducing the expression of IKZF3 (Aiolos) which silences the expression of IL-2^74^. Here, we describe for the first time a direct interaction between IRF4 and AHR, with combined motif analysis additionally revealing that AHR and IRF4 binding motifs are in close proximity (less than 5 bp apart).

Other examples for which, to our knowledge, no IRF4 heterodimer binding was demonstrated before are EP300, FLI1 and TCF7. In our dataset, we found an IRF4-EP300 steered regulation of Th17 and iTreg effector functions supporting data from previous studies, which reported a role of EP300 in Treg proliferation and suppressive function as well as FOXP3 stability^75,76^. In case of Th17 cells, EP300 is essential for the expression of pro-inflammatory cytokines and Th17 cell fate^77^. FLI1, enriched in the Th17 interactome, has been previously associated with the development of pro-inflammatory diseases, such as graft-versus-host disease, colitis and lupus^78–82^. *Fli1^−/−^*studies demonstrated a down-regulation of genes associated with pro-inflammatory states, while genes with anti-inflammatory properties were up-regulated^79,80,82^. Additionally, metabolic pathways were also affected by FLI1 knockout in T cells, which displayed decreased glycolytic but increased oxidative phosphorylation activity^78^ in line with our data. TCF7 is essential for a normal chromatin landscape in T cells^83^, acting mainly as a positive regulator on chromatin accessibility^39^ and is critical for formation of long-term memory T cells^84^ and Th17 longevity^38^. Neighbouring IRF and TCF7 binding sites were described along with a co-occupancy of IRF4 and TCF7 for chromatin accessibility in T cells^85^. In CD8^+^ cytotoxic cells IRF4 is involved in the expression of molecules essential for effector cell differentiation, including TCF7^20^.

To conclude, in the present study we integrate IRF4-ChIP-Seq, IRF4 interactome and quantitative proteome analysis to investigate the underlying molecular mechanisms of IRF4-steered transcriptional regulation in effector Th17 and iTreg cells. While we could demonstrate that IRF4 plays a more prevalent role in Th17 cells, we also defined cell type-specific gene regulatory networks required for both, Th17 and iTreg functionality. Moreover, this is the first interactome dataset that characterizes lineage-specific, endogenous IRF4 interaction partners identifying novel interactors that have not yet been described in Th17/Treg context. In addition, by linking the information derived from interactome analysis with ChIP-Seq data, we identified novel composite sequences targeted by the IRF4 complex involved in the regulation of Th17-and iTreg-specific transcriptional programs. In conclusion, our comprehensive dataset provides a valuable resource for studying IRF4-mediated gene regulatory programs in pro- and anti-inflammatory immune responses.

## METHODS

Methods and any associated references are available in the online version of the paper.

## ACKNOWLEDGEMENTS

We thank Ruben Spohrer and Christina Jung for excellent technical assistance. This work was supported by the German Research Foundation (DFG; Project Number 318346496, SFB1292/2 TP01 to T.B., TP11 to U.D. & N.L., and TP-Q1 to S.T., as well as DI 2471/1-1 to U.D. and BO 3306/2-1 to T.B.). The work was further supported by DFG priority program SPP 2225 (Grant No. TE599/9-1 to S.T.) and the SFB TRR355 (TPA9 and TPA10 to T.B.) and the by the Research Center for Immunotherapy (FZI) of the Johannes Gutenberg-University Mainz.

## AUTHOR CONTRIBUTIONS

T.B. established the transgenic mouse model. A.G. and S.D. performed the animal experiments and conducted the wet lab work. A.G., U.D. and G.H. designed and conducted the microscopic experiments. B.W., K.P. and A.G. performed the flow cytometric analysis. U.D. performed the mass spectrometric analyses. A.G. and M.K. performed the ChIP-Seq analysis. A.G., M.S., M.C., M.K., D.G.Z, T.Z. and U.D. analyzed the mass spectrometric and ChIP-Seq data. M.S. and M.C. wrote the R scripts and codes for downstream data analysis and data visualization. N.L., S.B., and S.T. aided the conceptualization of some experiments and contributed to writing the manuscript. U.D. and T.B. designed the project. A.G., M.S., M.C. and U.D. prepared the initial draft of the manuscript. All authors reviewed the final version of the manuscript.

## ONLINE METHODS

### Reagents and antibodies

Unless otherwise stated, all solvents (HPLC and Ultra LC-MS grade) were purchased from Roth and all chemicals were obtained from Sigma.

The following antibodies were used for fluorescence-activated cell sorting (FACS) analysis of murine cells: brilliant violet 711-conjugated rat anti-mouse CD4 (clone GK1.5, BD Biosciences), phycoerythrin-conjugated rat anti-mouse CD62L (clone MEL-14, BioLegend), brilliant violet 605-conjugated rat anti-mouse/human CD44 (clone IM7, BioLegend), allophycocyanin-conjugated rat anti-mouse/human RORγt (clone AFKJS-9, Thermo Fisher Scientific), phycoerythrin-conjugated rat anti-mouse IL-17A (clone eBio17B7, Thermo Fisher Scientific), fluorescein isothiocyanate-conjugated rat anti-mouse CD25 (clone 3C7, BioLegend), phycoerythrin-conjugated rat anti-mouse FOXP3 (clone FJK-16s, eBiosciences), phycoerythrin- and allophycocyanin-conjugated rat IgG2a,κ isotype controls (Thermo Fisher Scientific).

Polyclonal rabbit anti-IRF4 (P173), horseradish peroxidase-conjugated anti-rabbit IgG antibodies as well as horseradish peroxidase-conjugated anti-mouse IgG for Western blot analysis were obtained from Cell Signaling Technology. Monoclonal mouse anti-IRF4 (F-4, clone sc-48338) was purchased from Santa Cruz Biotechnology.

For confocal microscopy analysis, the following antibodies were used: monoclonal phycoerythrin-conjugated rat anti-human/mouse IRF4 (clone 3E4, Thermo Fisher Scientific), phycoerythrin-conjugated rat IgG1κ isotype control (BioLegend) and allophycocyanin-conjugated rat anti-mouse/human CD4 (clone GK1.5, Thermo Fisher Scientific).

### Animals

Bacterial artificial chromosome (BAC) transgenic mice (C57BL/6J-*Irf4^em1Bopp^*, referred to as IRF4^Avi-tag^ mice in the manuscript) were ordered from Cyagen US Inc. (Santa Clara, CA, USA). Detailed information on the IRF4^Avi-tag^ mouse strain expressing a fusion protein of IRF4 and the BirA recognition site (Avi-tag) under the control of the endogenous *Irf4* promotor is provided in Supplementary Fig. 5. IRF4^Avi-tag^ mice were cross-bred with the ROSA26^BirA^ (*Gt(ROSA)26^Sortm1.1(birA)Mejr^*) mouse strain ubiquitously expressing BirA ligase under the Rosa26 promotor^16^. ROSA26^BirA^ mice were kindly provided by Frank Großfeld/Dr. S. Klein-Heßling (Marburg, Germany). As a result of cross-breeding IRF4^Avi-tag^ and ROSA26^BirA^ mice, IRF4^Bio^ mice (C57BL/6J-*Irf4^em1Bopp^ Gt(ROSA)26^Sortm1.1(birA)Mejr^*) express biotinylated IRF4 due to BirA-mediated *in vivo* biotinylation of the IRF4-Avi-tag fusion protein. IRF4^Bio^ animals were used to conduct IRF4 interactome, proteome as well as IRF4 ChIP-seq experiments. Proteomic analyses of *Irf4^−/−^* and littermate control animals were performed using CD4^Cre^xIRF4^fl/fl^ mice, which were generated by crossing CD4^Cre^ ^86^ with IRF4^fl/fl^ animals^87^.

All animals are based on C57BL/6 background and were bred and housed at the Translational Animal Research Center (TARC) of the Johannes Gutenberg University of Mainz (Mainz, Germany) according to institutionally approved protocols.

### Isolation of murine splenocytes, naïve CD4^+^ T cells and Th17/iTreg differentiation

In order to obtain splenocytes as well as naïve CD4^+^ T cells, mice were sacrificed by CO_2_ asphyxiation. Afterwards, spleens of IRF4^Bio^ and ROSA26^BirA^ (control) mice were isolated, mechanically disrupted, and filtered from cell debris through a 40-μm cell strainer resulting in splenic single-cell suspensions.

Naïve CD4^+^ T cells were isolated from splenic single-cell suspensions by negative selecting magnetic-activated cell sorting (MACS) using the Naïve CD4^+^ T Cell Isolation Kit (mouse, Miltenyi Biotec) according to the manufacturer’s protocol. In brief, cells were centrifuged and cell pellets resuspended in GM-buffer (0.5 % (w/v) bovine serum albumin (BSA), 5 mM EDTA in phosphate-buffered saline (PBS), pH 7.3). After addition of the biotin-conjugated monoclonal antibody cocktail provided in the Naïve CD4^+^ T Cell Isolation Kit, cells were incubated at 4 °C for 5 min. Afterwards, cells were further diluted with GM-buffer and anti-biotin microbeads as well as anti-CD44 microbeads were added to the samples, followed by incubation for 10 min at 4 °C. Magnetic cell separation was performed using LS columns (Miltenyi Biotec). After column equilibration, up to 1 × 10^8^ cells were loaded per column and columns were washed twice with GM-Buffer. The flow-through, containing unlabeled naïve CD4^+^ cells, was collected and centrifuged at 630 xg for 10 min at 4 °C. After centrifugation, naïve CD4^+^ cells were reconstituted in 10 % (v/v) FCS, 1 % (w/v) sodium pyruvate, 1 % (w/v) glutamine in Iscove’s Modified Dulbecco’s Medium (IMDM, PAN Biotech), pH 7.2 and transferred to 24-well cell culture plates, which were coated with anti-CD3 and anti-CD28 antibodies as described before^88^. In total, 1 × 10^6^ cells were seeded per well. After the addition of polarizing cytokines, cells were cultured for 72 h at 37 °C in a 5 % CO_2_ environment. Th17 cell differentiation was induced adding 1 ng/mL TGF-ß, 40 ng/mL IL-6, 2.5 µg/mL anti-IFN-y and 10 µg/mL anti-IL4 to the medium while for Treg cells 7.5 ng/ml TGF-ß and 250 ng/ml IL2 was added. For each experiment, splenocytes of three mice were pooled prior to isolation of naïve CD4^+^ T cells (to obtain sufficient material, i.e. 3-5 ×10^7^ Th17/Treg cells, after *ex vivo* differentiation for the AP-procedure).

### Flow cytometry

Purity of isolated, naïve CD4^+^ T and differentiated Th17/iTreg cells was analyzed by flow cytometry. Flow cytometry data were acquired on a BD FACSCanto flow cytometer and analyzed using FACSDiva (version 6.1.3, BD) as well as the FlowJo (version 10, BD) software packages.

To characterize naïve CD4^+^ T cells, cells were stained for 15 min at 4 °C with the eBioscience Fixable Viability Dye eFlour780 (Thermo Fisher Scientific) and labeled with antibodies against appropriate cell surface markers including brilliant violet 711-conjugated anti-CD4 (BD Bioscience), phycoerythrin-conjugated anti-CD62L (BioLegend) and brilliant violet 605-conjugated anti-CD44 (BioLegend). Cells were washed and resuspended in 0.5 % (w/v) BSA, 5 mM EDTA in PBS (GM-buffer) prior to FACS analysis.

Th17 differentiation was characterized by determining RORγt expression and IL-17 production. Cells were re-stimulated for 3 h at 37 °C in a 5 % CO_2_ atmosphere with phorbol myristate acetate (PMA), ionomycin and monensin. After re-stimulation, cells were stained with the eBioscience Fixable Viability Dye eFlour780 and brilliant violet 711-conjugated anti-CD4 as described above. For intracellular staining of RORγt and IL-17A, cells were fixed and permeabilized for 30 min at 4 °C using the Fix/Perm Buffer from Thermo Fisher Scientific. Afterwards, cells were stained with allophycocyanin-conjugated anti-RORγt (Thermo Fisher Scientific) and phycoerythrin-conjugated anti-IL-17A (Thermo Fisher Scientific) in Permeabilization Buffer (Thermo Fisher Scientific) for 30 min at 4 °C. Cells were washed and resuspended in 0.5 % (w/v) BSA, 5 mM EDTA in PBS (GM-buffer) prior to FACS analysis.

Treg differentiation was determined by the expression of CD25 (fluorescein isothiocyanate-conjugated anti-CD25, BioLegend) and FOXP3 (phycoerythrin-conjugated anti-FOXP3, eBioscience). Surface (viability, anti-CD4 and anti-CD25) as well as intracellular (anti-FOXP3) staining was performed as described above for Th17 cells.

### Chemical cross-linking and isolation of nuclei

Murine Th17/Treg cells were harvested and washed twice with PBS to remove remaining media. A fresh Dithiobis[succinimidyl propionate] (DSP, Thermo Fisher Scientific) solution (20 mM) in DMSO was prepared for each experiment and further diluted in PBS, pH 8.3 to a final working concentration of 0.75 mM DSP. Cells were cross-linked according to the manufacturer’s protocol for 30 min at room temperature. The reaction was stopped by adding Tris (pH 7.5) to a final concentration of 10 mM, followed by incubation for 15 min at room temperature. Afterwards, cells were washed twice with PBS to remove remaining DSP. In order to isolate cell nuclei, cells were resuspended in a hypotonic buffer (20 mM Tris-HCl, 10 mM NaCl, 3 mM MgCl_2_, pH 7.4). After incubation for 12 min on ice, NP-40 was added to a final concentration of 0.5 % (v/v) to the samples and cells were vortexed for 10 s to disrupt cell membranes. Nuclei were washed twice with PBS and pelleted by centrifugation at 850 xg for 10 min at 4 °C. Isolated nuclei were either stored at −80 °C until further processing or directly subjected to immunoprecipitation.

### Isolation of IRF4 complexes

Nuclei were resuspended in Cell Lysis Buffer from the µMACS FactorFinder Kit (Miltenyi Biotec), which was supplemented with proteinase inhibitor cocktail according to manufactureŕs instructions (Roche). Nuclear lysis and disruption of DNA was promoted by sonication at 4 °C for 10 min using a Bioruptor Plus (low, 30 s/30 s ON/OFF, Diagenode). After lysis of nuclei, samples were centrifuged for 6 min at 14,850 xg at 4 °C and supernatant containing nuclear proteins was subjected to immunoprecipitation. Streptavidin-coated magnetic beads (Dynabeads™ M-280, Thermo Fisher Scientific) were washed with a mixture of Cell Lysis and Binding Buffer (1:3 (v/v), µMACS FactorFinder Kit, Miltenyi Biotec) for 10 min at room temperature. Afterwards, beads were resuspended in Binding Buffer supplemented with Binding Enhancer (both derived from the µMACS FactorFinder Kit, Miltenyi Biotec) and added to the nuclear lysates. After overnight incubation at 4 °C, samples were incubated for 2 min on a magnetic rack until beads were settled. Supernatant was discarded and samples were washed three times with RIPA buffer (25 mM Tris-HCl, 150 mM NaCl, 1 % (v/v) NP-40, 1 % (w/v) sodium deoxycholate, 0.35 % (w/v) SDS, pH 7.5) for 10 min at room temperature. Afterwards, the samples were washed three times with 25 mM Tris (pH 7.5) for 10 min at room temperature. Proteins were eluted incubating the streptavidin-coated beads in an SDS/biotin-containing buffer (1 % (w/v) SDS, 10 mM biotin, 10 mM Tris pH 7.5) for 5 min at 95 °C.

### Proteolytic digestion

Proteins released from IRF4 pulldown experiments and murine Th17/Treg cell lysates were processed by single-pot solid-phase-enhanced sample preparation (SP3) as detailed before with minor modifications^89,90^: In case of IRF4 pulldown experiments, proteins and DSP cross-linker were reduced with 45 mM DTT for 30 min at 37 °C. Afterwards, temperature was increased to 45 °C and samples were incubated for another 10 minutes. Whole cellular lysates were incubated with 20 mM DTT for 30 min at 45 °C. In all cases (IRF4 pulldown and sample preparation of murine Th17/Treg cells), proteins were alkylated for 30 min at room temperature using iodoacetamide (IAA). Excess IAA was quenched by the addition of DTT. Afterwards, 2 µL of carboxylate-modified paramagnetic beads (Sera-Mag SpeedBeads, GE Healthcare, 0.5 μg solids/μL in water as described by Hughes *et al*.^89^) were added to the samples. After adding acetonitrile to a final concentration of 70 % (v/v), samples were allowed to settle at room temperature for 20 min. Samples were mixed after 10 min. Subsequently, beads were immobilized by incubation on a magnetic rack for 2 min and washed twice with 70 % (v/v) ethanol in water and once with acetonitrile. Beads were resuspended in 50 mM NH_4_HCO_3_ supplemented with trypsin (Mass Spectrometry Grade, Promega) at an enzyme-to-protein ratio of 1:25 (w/w) and incubated overnight at 37 °C. After overnight digestion, acetonitrile was added to the samples to reach a final concentration of 95 % (v/v). Subsequently, samples were incubated for 20 min at room temperature (and mixed after 10 min). To increase the yield, supernatants derived from this initial peptide-binding step were additionally subjected to the SP3 peptide purification procedure as described before^90^. Each sample was washed with acetonitrile. To recover bound peptides, paramagnetic beads from the original sample and corresponding supernatants were pooled in 2 % (v/v) dimethyl sulfoxide (DMSO) in water and sonicated for 1 min. After 2 min of centrifugation at 16,200xg and 4 °C, supernatants containing tryptic peptides were transferred into a glass vial for MS analysis and acidified with 0.1 % (v/v) formic acid.

### Western blot analysis

Protein samples were separated via SDS-Polyacrylamide Gel Electrophoresis (SDS-PAGE). After electrophoretic separation, proteins were transferred onto a PVFD membrane (0.45 µm pore size, GE Healthcare). The membrane was blocked with 5 % (w/v) BSA in TBS-T buffer (1 mM Tris-base, 9 mM Tris-HCl, 150 mM NaCl, 1 % (v/v) Tween in H_2_O) for 1 h, followed by incubation with a primary polyclonal rabbit anti-IRF4 antibody (P173, Cell Signaling Technology, 1:1,000) overnight. A secondary horseradish peroxidase– conjugated anti-rabbit IgG (Cell Signaling Technology, 1:2,000) was used for detection.

Biotinylated proteins were detected using a horseradish peroxidase–conjugated streptavidin (Roche, 1:1,000) and visualized using WesternBright Chemiluminescence substrates (Biozym). The chemiluminescence signal was captured with a Bio-Rad ChemiDoc XRS imager and analyzed using the Quantity One software (version 4.4.0, Bio-Rad).

### Liquid chromatography-mass spectrometry (LC-MS) analysis

LC-MS analyses from initial AP experiments during protocol optimization (i.e., splenocytes) were conducted on a nanoAcquity UPLC system (Waters Corporation), which was coupled online to a Synapt G2-S HDMS mass spectrometer (Waters Corporation) via a NanoLockSpray dual electrospray ion source (Waters Corporation). Tryptic digests were loaded onto an HSS-T3 C18 1.8 μm, 75 μm × 250 analytical reversed-phase column (Waters Corporation) in direct injection mode as described before^91^. Mobile phase A was water containing 0.1 % v/v formic acid and 3 % (v/v) DMSO, while mobile phase B was ACN containing 0.1 % v/v formic acid and 3 % (v/v) DMSO in ACN^92^. Peptides were separated running a gradient of 5–40 % mobile phase B over 90 min at a flow rate of 300 nL/min. Afterwards, the column was rinsed for 5 min with 90 % mobile phase B, followed by a re-equilibration step at initial conditions resulting in a total analysis time of 120 min. The column temperature was maintained at 55 °C. MS analysis of eluting peptides was performed by ion-mobility separation (IMS) enhanced data-independent acquisition (DIA) in UDMS^E^ mode as detailed before^91,93^.

LC-MS analyses of murine Th17/iTreg IRF4 interactome and proteome were performed on an Ultimate 3000 RSLCnano LC system (Thermo Fisher Scientific) coupled to an Orbitrap Exploris 480 instrument platform (Thermo Fisher Scientific). Tryptic peptides were first loaded onto a PEPMAP100 C18 5 µm 0.3 × 5 mm trap column (Thermo Fisher Scientific) and subsequently separated on an HSS-T3 C18 1.8 μm, 75 μm × 250 mm analytical reversed-phase column (Waters Corporation). Mobile phase A was water containing 0.1 % (v/v) formic acid and 3 % (v/v) DMSO. Peptides were separated running a gradient of 2–35 % mobile phase B (0.1 % (v/v) formic acid, 3 % (v/v) DMSO in ACN) over 40 min at a flow rate of 300 nL/min. Total analysis time was 60 min including wash and column re-equilibration steps. Column temperature was set to 55 °C. The following settings were used for mass spectrometric analysis of eluting peptides on the Orbitrap Exploris 480 instrument platform: Spray voltage was set to 1.8 kV, the funnel RF level to 40, and heated capillary temperature was at 275 °C. Data were acquired in DIA mode. Full MS resolution was set to 120,000 at *m/z* 200 and full MS automated gain control (AGC) target to 300 % with a maximum injection time (IT) of 20 ms. Mass range was set to *m/z* 345 – 1250. Fragment ion spectra were acquired with an AGC target value of 1000 %. In total, 20 windows with varying sizes (adjusted to precursor density) were used with an overlap of 0.5 Th. Resolution was set to 30,000 and IT was determined automatically (“auto mode”). Normalized collision energy was fixed at 27 %. All data were acquired in profile mode using positive polarity.

For the analysis of the IRF4 knock-out data set, reconstituted peptides were injected and separated on a nanoElute LC system (Bruker Corporation, Billerica, MA, USA) at 400 nL/min using a reversed phase C18 column (Aurora UHPLC emitter column, 25 cm × 75 µm 1.6 µm, IonOpticks) which was heated to 50°C. Peptides were loaded onto the column in direct injection mode at 600 bar. Mobile phase A was 0.1% FA (v/v) in water and mobile phase B 0.1% FA (v/v) in ACN. Peptides were separated running a linear gradient from 2% to 37% mobile phase B over 39 min. Afterwards, column was rinsed for 5 min at 95% B. Eluting peptides were analyzed in positive mode ESI-MS using parallel accumulation serial fragmentation (PASEF) enhanced DIA mode on a timsTOF Pro 2 mass spectrometer (Bruker Corporation)^94^. The dual TIMS (trapped ion mobility spectrometer) was operated at a fixed duty cycle close to 100% using equal accumulation and ramp times of 100Lms each spanning a mobility range from 1/K_0_L=L0.6LVsLcm^−2^ to 1.6LVsLcm^−2^. We defined 36L×L25LTh isolation windows from *m/z* 300 to 1,165 resulting in 15 diaPASEF scans per acquisition cycle. The collision energy was ramped linearly as a function of the mobility from 59LeV at 1/K_0_L=L1.3LVsLcm^−2^ to 20LeV at 1/K_0_L=L0.85LVsLcm^−2^.

All samples were analyzed in three technical replicates.

### Data analysis and label-free quantification

Processing and database search of the Synapt G2-S raw data were performed using ProteinLynx Global SERVER (PLGS, version 3.0.2, Waters Corporation). Data were searched against a custom compiled database containing UniProtKB/SwissProt entries of the mouse reference proteome (UniProtKB release 2020_03, 17,033 entries) and a list of common contaminants as described before^91^. Postprocessing of data including retention time alignment, EMRT (exact mass retention time) and IMS clustering, as well as protein homology filtering was conducted using the software tool ISOQuant (ver.1.8) as detailed before^91,93^. Each pulldown experiment was processed separately. An experiment-wide

FDR of 0.01 was applied at the peptide-level for cluster annotation in ISOQuant. Moreover, only proteins that had been identified by at least two peptides with a minimum length of 7 amino acids, a minimum PLGS score of 6.0 and no missed cleavages were considered for quantitative analysis in the final dataset. For each protein absolute in-sample amounts were calculated using TOP3 quantification as described before^95^. IRF4 interacting proteins had to show a 2-fold enrichment as compared to the controls.

DIA raw data acquired with the Exploris 480 and the timsTOF Pro2 platforms were processed using DIA-NN (version 1.7.13)^96^ applying the default parameters for library-free database search. Data were searched using a custom compiled database containing UniProtKB/SwissProt entries of the murine reference proteome (UniProtKB release 2021_04, 17,068 entries) and a list of common contaminants. For peptide identification and *in-silico* library generation, trypsin was set as protease allowing one missed cleavage. Carbamidomethylation was set as fixed modification and the maximum number of variable modifications was set to zero. The peptide length ranged between 7–30 amino acids. The precursor *m/z* range was set to 300–1,800, and the product ion *m/z* range to 200–1,800. As quantification strategy we applied the “any LC (high accuracy)” mode with RT-dependent median-based cross-run normalization enabled. We used the build-in algorithm of DIA-NN to automatically optimize MS2 and MS1 mass accuracies and scan window size. Peptide precursor FDRs were controlled below 1 %. In the final proteome datasets, proteins had to be identified by at least two peptides and had to be present in at least eight runs in at least one of the conditions. In the proteome analysis of Th17/iTreg cells from *Irf4^−/−^*and littermate controls, the run filter was set to at least seven runs out of twelve in at least one condition. The final IRF4 interactome dataset was filtered to only contain proteins identified by more than two peptides and present in at least eight out of nine runs in at least one of the conditions. Additionally, the cross-run normalization was disabled.

### Statistical Analysis of proteomic datasets

Statistical Analysis was conducted with the limma package^97^. For the interactome data, proteins, identified in at least eight out of nine LC-MS runs in the IRF4^Bio^ condition, with an adjusted *p*-value of 0.01 and a log_2_-fold change (IRF4^Bio^/Ctrl) larger than 1 were considered as significantly enriched. In case of the proteomic datasets, proteins with an adjusted *p*-value of 0.01 and a log_2_-fold change either below −0.5 or above 0.5 were considered significant differentially expressed.

To enable limma to work with the intensities, they were log transformed. We utilized DEPs mixed imputation method to impute missing data differently for missing at random (MAR) and missing not at random (MNAR) data: MNAR values were imputed with the minimum value present in the data, while MAR values were imputed with K-Nearest Neighbor imputation, both as implemented in the DEP package^98^. Despite limma’s capabilities of working with data with missing values, we favored imputation as some of the most interesting proteins, would be those that could not be identified or could not pass the detection threshold in the control, but would do so in the IRF4^Bio^ data. For the proteome, in both cell types compared, proteins that passed the threshold in only one of the groups would be of interest.

The design for limma’s linear model was different for the proteome and the interactome analysis respectively: For the proteome analysis just the cell type, which was the comparison of interest, was given to the model as factor. In the interactome analysis, additional to the comparison of interest, which was the control mice against the IRF4^Bio^ mice, we added the experiments as a co-factor. This was done to counteract the bias of an outlier experiment, which had generally lower intensities than its partner experiments.

Principal component analysis was conducted for plotting using the package FactoMineR^99^ and heatmaps were plotted with pheatmap^100^. ggplot2 was used for scatterplots^101^. Differential proteins or clusters in heatmaps were annotated with the GO terms with the clusterprofiler package^102^.

### Functional annotation and downstream data analysis

Functional annotation analysis was performed using GO terms powered by PANTHER (version 14.0)^103^ (http://geneontology.org/). Network analysis was conducted using the STRING database (version 11.0)^104^ through its web interface as well as the stringApp^105^ in Cytoscape (version 3.8.0)^106^. Protein networks in STRING were generated using default settings. Venn diagram data were calculated using the Venny web application (http://bioinfogp.cnb.csic.es/tools/venny/index.html).

### Chromatin Immunoprecipitation coupled with Sequencing (ChIP-seq)

For ChIP-seq experiments 1 – 1.5 × 10^7^ differentiated Th17 or iTreg cells were washed twice with medium containing 5 % (v/v) FCS, 1 % (w/v) sodium pyruvate, 1 % (w/v) glutamine in Iscove’s Modified Dulbecco’s Medium (IMDM, PAN Biotech), pH 7.2. The cells were fixed with a final concentration of 1 % formaldehyde (Pierce 16 % formaldehyde, methanol-free, ThermoFisher) for 7 min at room temperature. To quench the reaction a final concentration of 125 mM of glycerin was added to the solution for 5 min at room temperature. During the fixing and quenching procedure, the samples were inverted from time to time. To remove remaining medium components, the cells were washed twice with cold PBS and were transferred into TPX microtubes (Diagenode). The dry cell pellet was stored over night at –80 °C. Before adding the lysis buffer (50 mM Tris pH 7.9, 10 mM EDTA pH 8, 1 % (w/v) SDS in nuclease-free water, QIAGEN) supplemented with proteinase inhibitor cocktail (Roche), the cell pellet was scratched over a rack to loosen it. The lysis buffer was incubated at least for 30 min on ice followed by the sheering of the cells 6 times with a 1 mL syringe through a 21G × 1^1/2^ ’’ 0.8 × 40 mm needle on ice. To fragment the chromatin for an average size of 200-500 bp, the cells were sonicated 3-6 times (cycles) at 4 °C each for 10 min using the Bioruptor Plus (high, 30 s/30 s ON/OFF, Diagenode). Between every cycle, the cells were inverted and quickly spun down. The supernatant was collected in a new 1.5 mL tube after centrifugation 15 min at 17,000 xg, 4 °C. Afterwards the DNA content was determined and adjusted between the IRF4^Bio^ and ROSA26^BirA^ (control) sample. Streptavidin-coated magnetic beads (Dynabeads™ M-280, Thermo Fisher Scientific) were equilibrated with 80 % (v/v) IP buffer (30 mM Tris pH 7.9, 2 mM EDTA pH 8, 165 mM NaCl, 0.3 % (w/v) SDS, 1 % (v/v) Triton X-100 in nuclease-free water) and 20 % (v/v) of lysis buffer by rolling for 5-10 min at room temperature. IP buffer was added to the sample to gain a 4:1 ratio of IP:lysis buffer. A small amount of sample was removed serving as an input control in the real-time PCR after chromatin immunoprecipitation. The washed streptavidin-coated magnetic beads were added to the sample and incubated rolling for 4 h at room temperature before overnight incubation at 4 °C. The beads were washed once with IP buffer supplemented with proteinase inhibitor cocktail rolling 30 min at room temperature. After the beads settled on a magnetic rack the supernatant was discarded. Further washing steps followed, each step rolling for 30 min at room temperature and supplemented with proteinase inhibitor cocktail: twice with a washing buffer containing 2 % SDS (in nuclease-free water), twice with a washing buffer containing 10 mM Tris pH 7.9, 1 mM EDTA pH 8, 250 mM LiCl, 1 % (v/v) NP-40, 1 % (v/v) sodium deoxycholate (in nuclease-free water) and twice with a washing buffer containing 20 mM Tris pH 7.9, 1 mM ETDA pH 8, 50 mM EDTA, 0.1 % (v/v) SDS (in nucleas-free water). Before the addition of the elution buffer (10 mM Tris pH 7.9, 5 mM EDTA pH 8.0, 1 % (w/v) SDS, 500 mM NaCl in nuclease-free water), a 15 min washing step at room temperature only with nuclease-free water was included. For the treatment with RNAseA and proteinase K, the input control from the previous day was included and to all samples 50 µg/mL RNaseA (DNase and protease-free, ThemoFisher) was added to all samples. Samples were then incubated for 1 h at 37 °C. Afterwards, 150 µg/mL proteinase K (from *Tritirachium album*, Serva) was added and samples were incubated 90 min at 42 °C. All samples were incubating rolling at 65 °C overnight. The DNA Clean & Concentrator Kit (Zymo Research) was used to clean and concentrate the chromatin fragments according to the manufacturer’s protocol. Before the samples were sent for sequencing the fold enrichment of the immunoprecipitated chromatin was calculated using a real-time PCR. For that a target region, where IRF4 binds (IL10, fwd: 5’AATCCGAGAAACCCACCA3’, rev: 5’ TCCATACCAAAACCCCAG3’) as well as an off-target region, where IRF4 does not bind (Ezh2, fwd: 5’CTTCTCAACCCCTTTCCCTAAGA3’, rev: 5’CACCTTATTCCCAAAGGCAAGG3’) were selected. All samples with an at least 4-fold enrichment were used to generate a DNA library. For this the NEBNext Ultra II DNA Library Prep Kit for Illumina (New England BioLabs) was used according to the recommended protocol for low input material.

### Data analysis of sequencing data (ChIP-Seq)

We used seqQscorer^107^ for quality control of the raw data. For the downstream analysis of the sequencing data, the software EaSeq (version 1.2, https://easeq.net)^108^ was used to call peaks. As a reference, the mm10 genome was used. For peak calling in Th17 cells the following parameters were used: window size: 100 bp, FDR: 1×10^−15^, p-value: 1×10^−20^, log_2_: 2.1 and merge distance: 0 bp. For Treg cells the parameters were adjusted to: window size: 100 bp, FDR: 1×10^−10^, p-value: 1×10^−10^, log_2_: 1.5 and merge distance: 100 bp. The peaks were annotated to the nearest start or end of a gene (annotation “Start&End”). ChIPseeker^109^ was used to additionally annotate called peaks.

To search for enrichment in enhancer and silencer regions, we conducted a search in the silencers found in mouse blood according to silencerDB^48^ and the enhancers provided in the EnhancerAtlas 2.0^47^ and filtered for CD4^+^ cells as a source. As both these databases were using mm9 we used UCSCs LiftOver (https://genome.ucsc.edu/cgi-bin/hgLiftOver) to remap them to mm10. We used ChIPPeakAnno^110^ to search for an overlap between the two respective databases and the peaks of both Th17 and Treg cells. To ensure the found overlap is not random we conducted a random permutation test with 1,100 iterations as implemented in ChIPPeakAnno.

*Ab initio* motif enrichment analysis of the IRF4 ChIP-Seq peaks derived from Th17 and Treg cells was executed using the MEME suite with default parameters^52^. Identified motifs were annotated using Tomtom^53^ with a similarity threshold of 0.05, in conjunction with the position-specific scoring matrices (PSSM) of murine TFs from the TRANSFAC database^50^.

To enhance motif annotation, we additionally employed the MATCH algorithm^49^ annotating known TF binding sites. This process incorporated PSSMs corresponding to Th17 and Treg interactors, as well as the profile of immune-specific TFs as defined by TRANSFAC^50^. We established a stringent cutoff threshold, accepting motif annotations only if they matched both the core sequence and matrix sequence of PSSMs with a similarity score of 0.8 or higher.

To further investigate the combinatorial activity of TFs, we quantified the number of peaks bound by two distinct TF motifs within a spatial proximity of equal or less than five base pairs. This approach facilitated a detailed exploration into the complex interplay of TFs within Th17 and Treg cells.

### Immunostaining for Confocal Fluorescence Microscopy

Between 5×10^5^ to 1.5×10^6^ cells were used per preparation. After harvest, cells were washed twice with PBS or PBS supplemented with 0.5 % (w/v) BSA and 5 mM EDTA and subjected to immunostaining. Cells were first incubated with an allophycocyanin-conjugated anti-CD4 antibody (Thermo Fisher Scientific, 1:500 dilution) in PBS for 20 min at 4 °C. Afterwards, the cells were washed twice with PBS. Cells were subsequently fixed and permeabilized for 30 min at 4 °C using Fix/Perm Buffer (Thermo Fisher Scientific). Afterwards, cells were washed twice with Permeabilization Buffer (Thermo Fisher Scientific) and cells were incubated with phycoerythrin-conjugated anti-IRF4 antibody (Thermo Fisher Scientific, 1:500 dilution) in PBS for 30 min at 4 °C. After washing cells twice with PBS, cells were incubated with Hoechst (ImmunoChemistry Technologies, 1:500 dilution) in PBS for 5 min at 4 °C. Afterwards, cells were washed twice with PBS and resuspended in 40 µL of warm (37 °C) Mowiol (produced in-house). Reconstituted cells were then transferred onto a glass slide (Diagonal), covered with a no.1.5H cover glass (Paul Marienfeld GmbH) and samples were allowed to dry for 6 to 12 h at room temperature prior to storage at 4 °C and/or imaging at the confocal microscope.

### Confocal Microscopy

The fixed and stained cells were imaged on a Leica SP8 confocal microscope with either a 20X 0.75 NA air objective or with a 63X 1.40 oil immersion objective sequentially with exposure to a 405 nm laser for transmission images and for Hoechst excitation with emission and detection within a spectral window of 440 nm to 480 nm, with 552 nm laser exposure for phycoerythrin excitation and with emission and detection within a spectral window of 565 nm to 625 nm and with 638 nm laser exposure for allophycocyanin excitation with emission and detection within a spectral window of 650 nm to 735 nm. The images were always acquired at a factor of 2.3 times greater than the calculated confocal spatial lateral resolution, at a scan rate of 200 lines/second and with 4x averaging. All images used in comparison were prepared and acquired under the same conditions.

The images shown in the figures were smoothed with the standard Leica smoothing algorithm. In some cases, the Hoechst signal was processed by having the upper threshold lowered by up to 20 % in order to create homogenous nuclear images and homogenous cell nuclear borders. In some cases, the CD4 allophycocyanin signal had up to a 10 % cutoff and/or a 10 % threshold applied in order to create a homogeneous cell border. The IRF4 phycoerythrin signal intensity was left unchanged.

### Data availability

The mass spectrometry proteomics data have been deposited to the ProteomeXchange Consortium (http://proteomecentral.proteomexchange.org) via the PRIDE partner repository^111^ with the dataset identifier PXD044298. The ChIP-seq data can be accessed on gene expression omnibus (GEO) via the accession number GSE240979.

The R-scripts are made available at: https://github.com/Muedi/IRF4-in-Treg-and-Th17.

