## Supplementary Figures for "Unveiling IRF4-steered regulation of context-dependent effector programs in Th17 and Treg cells"

**Supplementary material:**


| | |
|---|---|
| **Supplementary Fig. 1** | String Network analysis of proteins associated with cluster 1 “signal receptor activity” displayed in Fig. 1e. |
| **Supplementary Fig. 2** | String Network analysis of proteins associated with cluster 3 “signal receptor regulator activity” displayed in Fig. 1e. |
| **Supplementary Fig. 3** | Proteome analysis of Th17 and iTreg cells differentiated from naïve $CD4^+$ T cells derived from WT and *Irf4*$^{-/-}$ mice reveals marked protein expression changes. |
| **Supplementary Fig. 4** | Proteome analysis of Th17 and iTreg cells differentiated from naïve $CD4^+$ T cells derived from WT and *Irf4*$^{-/-}$ mice reveals marked changes in pathways required for cellular effector functions. |
| **Supplementary Fig. 5** | Generation of IRF4$^{Avi\text{-}tag}$ animals. |
| **Supplementary Fig. 6** | Western blot analyses of IRF4$^{Bio}$ and ROSA26$^{BirA}$ (Ctrl) mice (non-cropped blots, data Fig. 2C). |
| **Supplementary Fig. 7** | Localization and expression of IRF4 three days after isolation of naïve $CD4^+$ T cells and Th17 *ex vivo* differentiation. |
| **Supplementary Fig. 8** | Exemplified gating strategies for FACS analysis and characterization of murine Th17 cells. |
| **Supplementary Fig. 9** | Exemplified gating strategies for FACS analysis and characterization of murine iTreg cells. |
| **Supplementary Fig. 10** | Optimization of the IRF4$^{Bio}$ affinity purification (AP) protocol. |
| **Supplementary Fig. 11** | FACS analysis of splenocytes and characterization of IRF4 positive subpopulations. |
| **Supplementary Fig. 12** | Optimized AP-MS protocol allows the *ex vivo* characterization of transcription factor interactomes from different subcellular localizations. |
| **Supplementary Fig. 13** | Nuclear IRF4 interactome in fully differentiated Th17 and iTreg cells. |
| **Supplementary Fig. 14** | Pathway analysis of proteins from the “IRF4 core” interactome reveals shared features between Th17 and iTreg cells. |
| **Supplementary Fig. 15** | Integrating proteome and ChIP-Seq analysis for the detection of proteins transcriptionally regulated by IRF4 which display aberrant expression levels in fully differentiated cells. |
| **Supplementary Fig. 16** | Pathway enrichment analysis of proteins that are regulated by IRF4 on DNA and protein level reveals cell type-specific functions. |

**Supplementary Fig. 1**

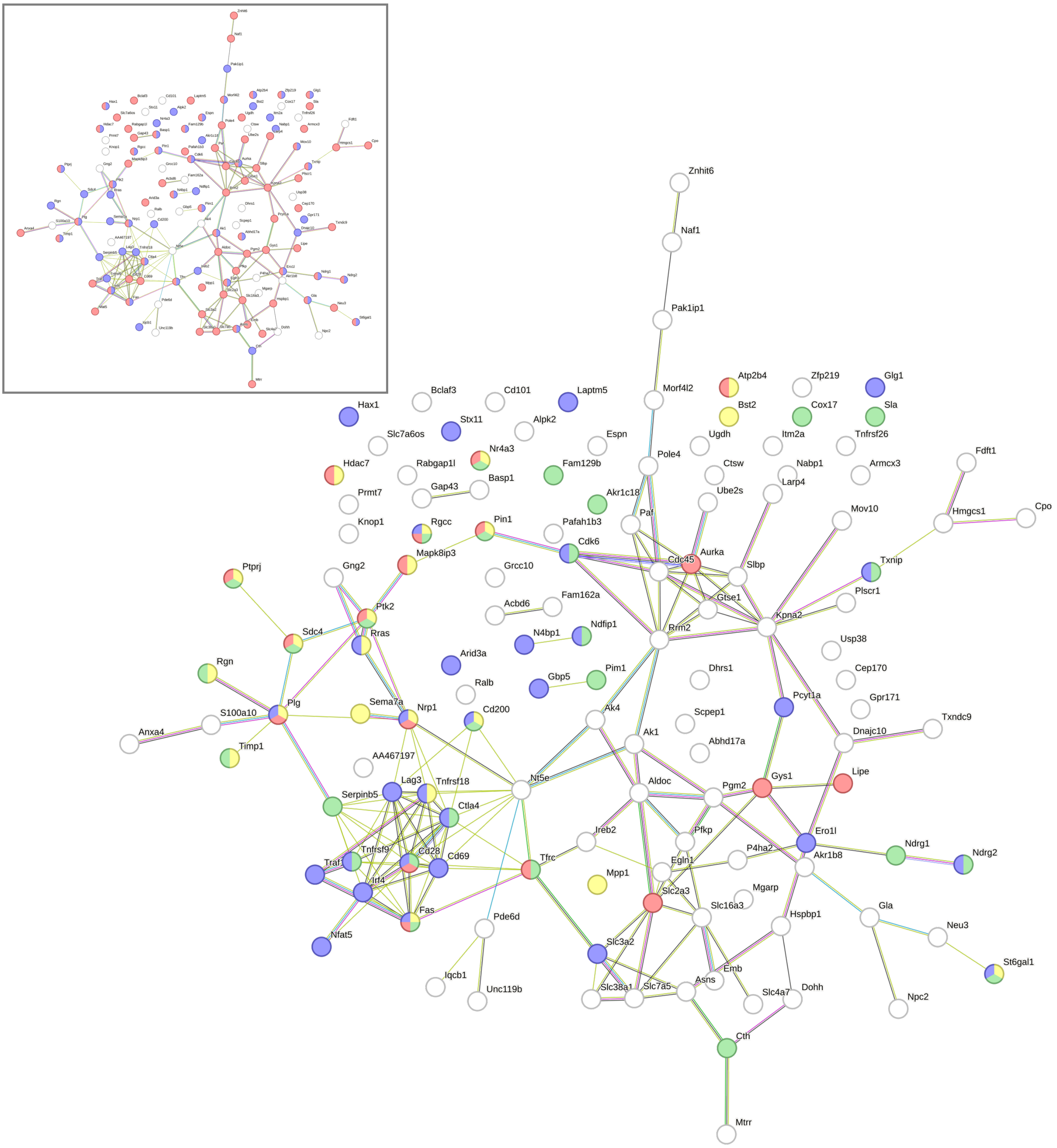


**String Network analysis of proteins associated with cluster 1 “signal receptor activity” displayed in Fig. 1e**. String network analysis revealed a strong enrichment of phosphoproteins (red, 85 out of 134 proteins) and proteins associated with the negative regulation of cellular processes (blue, 57 proteins) in cluster 1 (little inset). Within this cluster, we find multiple proteins associated with abnormal cell-mediated immunity (blue, including IRF4), kinase binding (red) and protein binding (68 proteins, not marked), as well as the regulation of locomotion (yellow) and cell population proliferation (green).

**Supplementary Fig. 2**

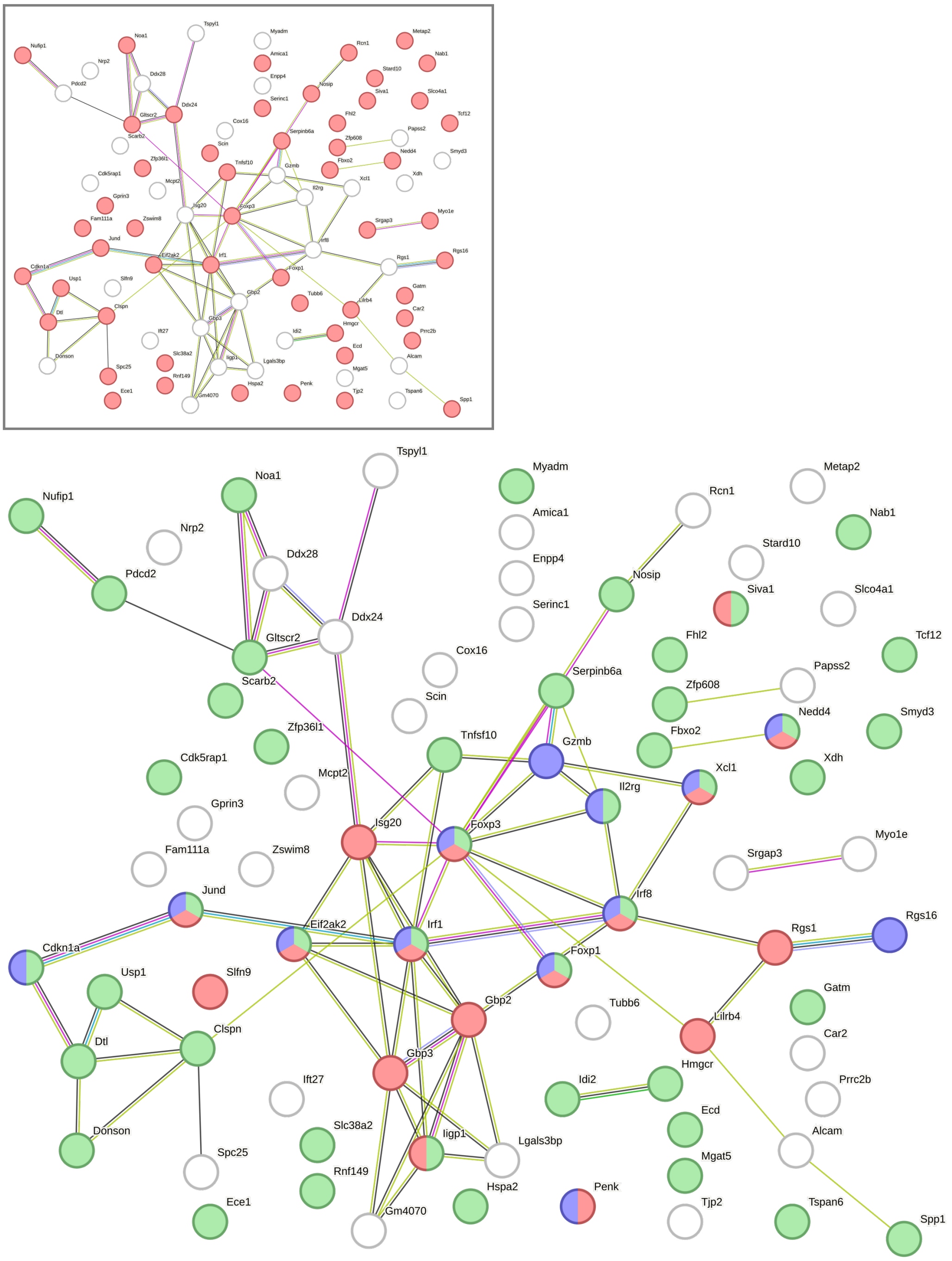


**String Network analysis of proteins associated with cluster 3 “signal receptor regulator activity” displayed in Fig. 1e.** Analogue to cluster 1 String network analysis revealed a strong enrichment of phosphoproteins (red, 52 out of 83 proteins) in cluster 3 (little inset). Within this cluster, we find a strong enrichment of proteins associated with the regulation of metabolic processes (green) and abnormal T cell physiology (blue) including various transcription factors, such as IRF1, IRF8, FOXP3, FOXP1, JunD, as well as proteins involved in interspecies interaction between organisms (red).

### Supplementary Fig. 3

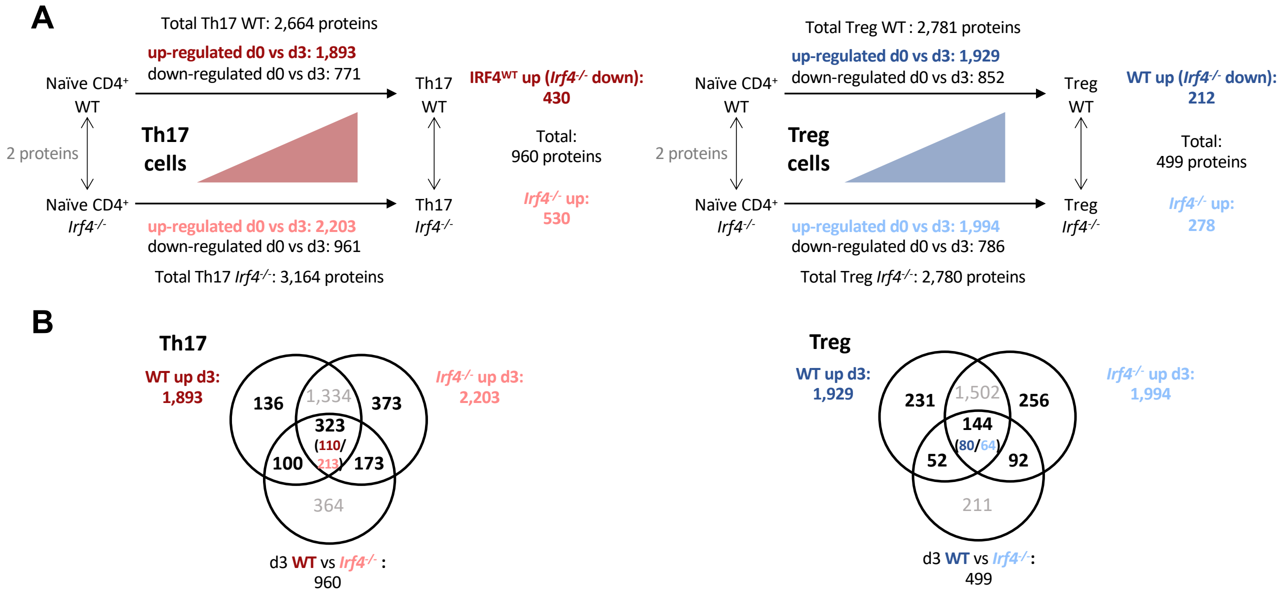


**Proteome analysis of Th17 and iTreg cells differentiated from naïve CD4$^{+}$ T cells derived from WT and *Irf4*$^{-/-}$ mice reveals marked protein expression changes**. (A) Numbers of differentially regulated proteins between naïve CD4$^{+}$ T (0h/d0) cells and fully differentiated Th17 and iTreg cells (72h/d3) and *Irf4*$^{-/-}$ as well as WT mice. Numbers next to the arrows marked as "total" indicate the overall number of differentially expressed proteins (including up- and downregulated proteins) between the different conditions (as displayed also in Fig. 1g). Numbers marked in bold indicate the number of up-regulated proteins between the different conditions (i.e., in the condition indicated) (B) Venn overlap of proteins that are up-regulated in *Irf4*$^{-/-}$ and WT animals upon differentiation (between 0h and 72h, marked in bold in (A)), with differentially regulated proteins at timepoint 72h. Proteins that show IRF4-dependencies marked in bold in the Venn diagram were used for Reactome pathway analysis as displayed in Fig. 1h.

### Supplementary Fig. 4

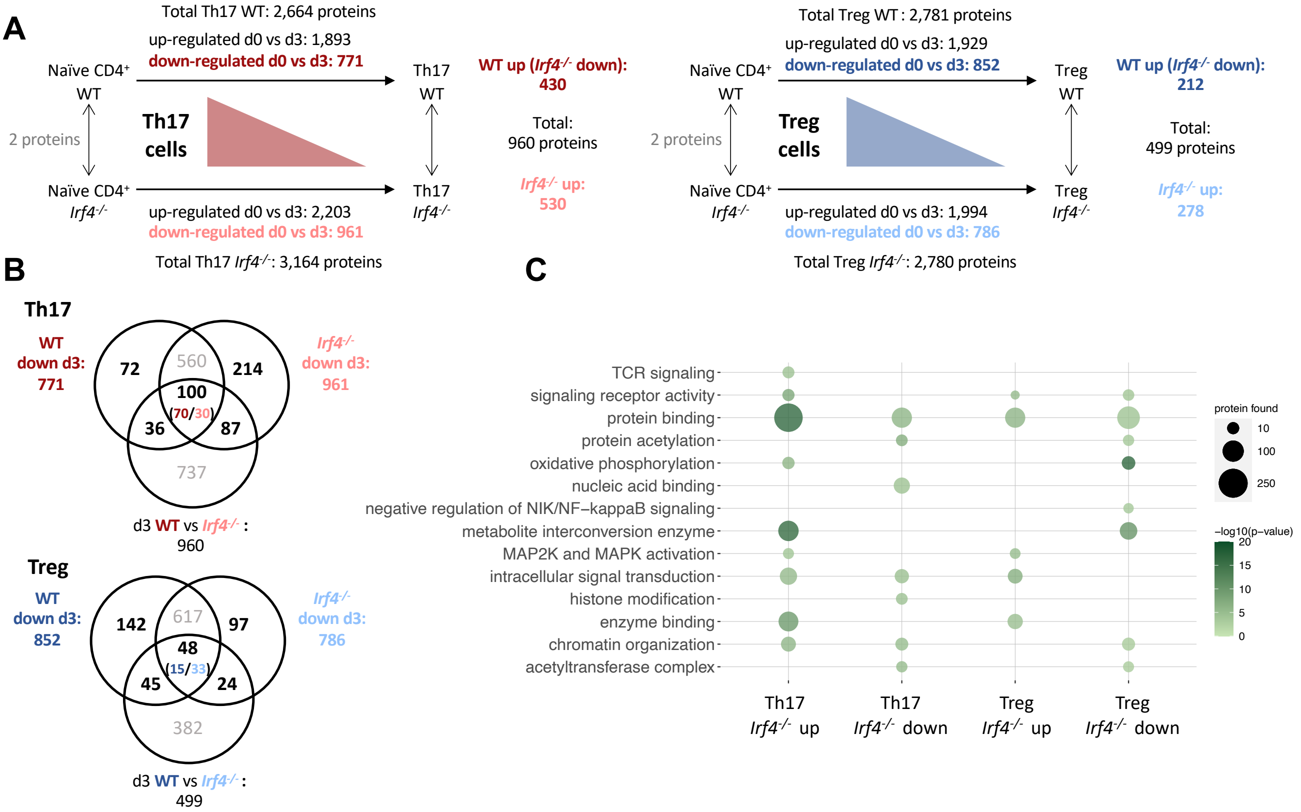


**Proteome analysis of Th17 and iTreg cells differentiated from naïve CD4$^{+}$ T cells derived from WT and *Irf4$^{-/-}$* mice reveals marked changes in pathways required for cellular effector functions**. (A) Numbers of differentially regulated proteins between naïve CD4$^{+}$ T (0h/d0) cells and fully differentiated Th17 and iTreg cells (72h/d3) and *Irf4$^{-/-}$* as well as WT mice. Numbers next to the arrows marked as "total" indicate the overall number of differentially expressed proteins (inclusive up- and downregulated proteins) between the different conditions. Numbers marked in bold indicate the number of down-regulated proteins between the different conditions (i.e., in the condition indicated). (B) Venn overlap of proteins that are down-regulated in *Irf4$^{-/-}$* and WT animals upon differentiation (between 0h and 72h, marked in bold in (A)), with differentially regulated proteins at timepoint 72h. Proteins that show IRF4-dependencies marked in bold in the Venn diagram were used for reactome pathway analysis. (C) Reactome pathway analysis of proteins that were downregulated on day three (72h) as compared to day zero (0h) but showed different behavior in Th17 and iTreg cells from WT and *Irf4$^{-/-}$* mice.

**Supplementary Fig. 5**

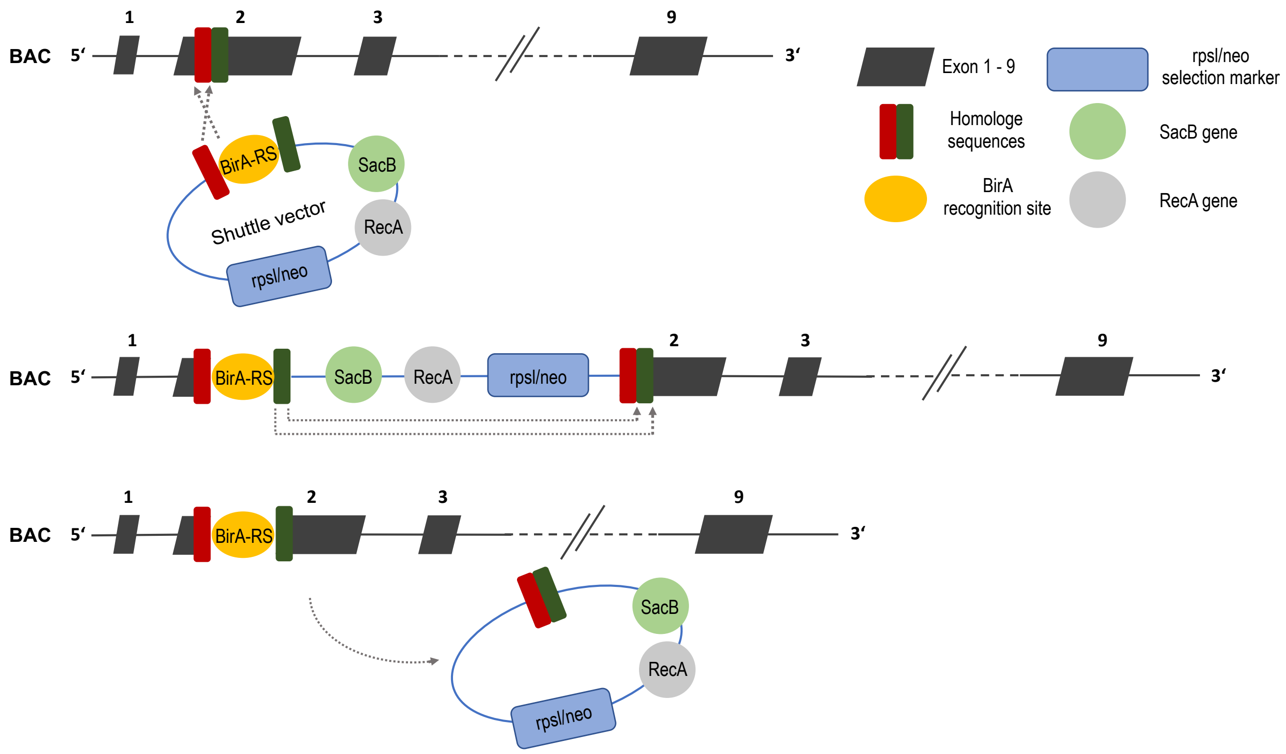


**Generation of IRF4$^{Avi\text{-}tag}$ animals.** In cooperation with Cyagen US Inc. (Santa Clara, CA, USA) a new bacterial artificial chromosome (BAC) transgenic mouse, C57BL/6J-*Irf4*$^{em1Bopp}$ referred to as IRF4$^{Avi\text{-}tag}$ mouse, was generated. IRF4$^{Avi\text{-}tag}$ transgenic mice code an IRF4-fusion protein located on a BAC transgene containing the *Irf4* sequence and the sequence for an *N*-terminal BirA recognition site (BirA-RS/Avi-tag) under the control of the endogenous *Irf4* promotor. To generate the new mouse strain, the sequence of the BirA recognition site was flanked with homologous sequences of a BAC (RP23-206G12) containing chromosome 13 with the *Irf4* gene by PCR and integrated into a shuttle vector containing *recA*, *sacB* (encoding levansucrase) and a rpsL-neo cassette (upper panel). Via RecA-mediated homologous recombination the shuttle vector was integrated in the BAC carrying the *Irf4* gene (exon 2, co-integration) in *E. coli*. After positive selection (i.e., resistance to neomycin), shuttle vector sequences were removed by a second homologous recombination (resolution, middle panel) resulting in a modified BAC transgene carrying sequences for the BirA recognition site (lower panel). After negative selection of successful resolution processes (i.e., loss of s*acB*), *Irf4*$^{Avi\text{-}tag}$-BAC was isolated and injected into murine blastocysts, which were transplanted into surrogate mothers. Successful integration of the BAC transgene was tested by PCR. Cross-breeding ROSA26$^{BirA}$ mice with IRF4$^{Avi\text{-}tag}$ animals results in a mouse strain, C57BL/6J-*Irf4*$^{em1Bopp}$ *Gt(ROSA)26*$^{Sortm1.1(birA)Mejr}$, that expresses biotinylated IRF4 (as demonstrated by Western blot analysis, see Fig. 2c).

**Supplementary Fig. 6**

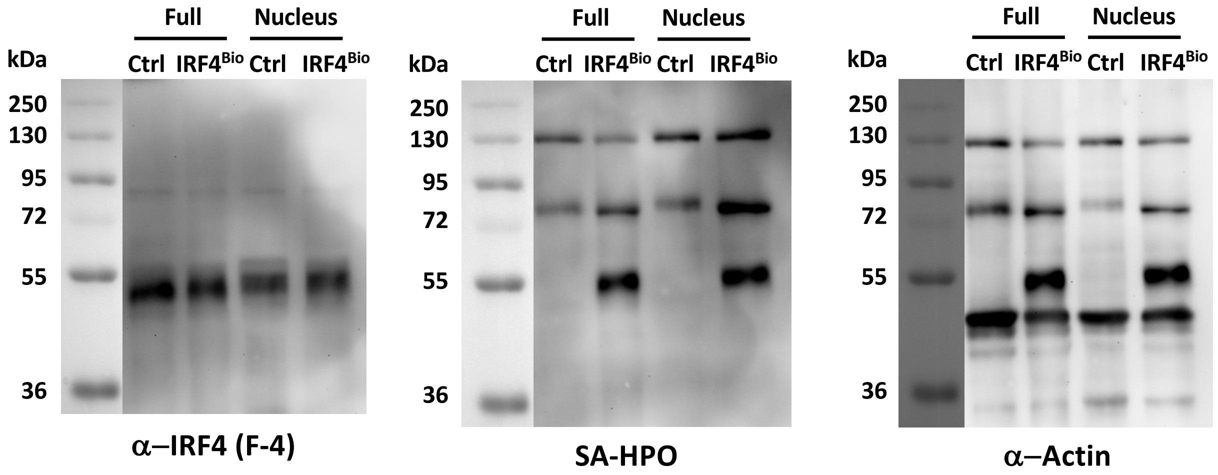


**Western blot analyses of IRF4$^{Bio}$ and ROSA26$^{BirA}$ (Ctrl) mice (non-cropped blots, data Fig. 2c)**. Full lysates as well as nuclear lysates of CD4$^{+}$ T cells (15 µg of total protein) were loaded on an SDS polyacrylamide gel. After gel electrophoresis, samples were analyzed by immunoblotting using a monoclonal mouse anti-IRF4 antibody (F-4, clone sc-48338, Santa Cruz Biotechnology, 1:500), a monoclonal mouse anti-actin HPO-conjugated antibody (clone AC-15, Sigma Life Science, 1:25,000) and horseradish peroxidase (HPO)–conjugated streptavidin (Roche, 1:1,000) to detect biotinylated proteins. After the detection of IRF4 (using an HPO-linked anti-mouse IgG secondary antibody, 1:2,000) and prior to incubation with HPO-conjugated streptavidin and the anti-actin antibody, membranes were stripped for 45 min at room temperature applying 0.1 M Glycin, 0.2 % (w/v) SDS in water (pH 2.5). Proteins were visualized using WesternBright Chemiluminescence substrates (Biozym) according to the manufacturer´s instructions. The chemiluminescence signal was captured with a Bio-Rad ChemiDoc XRS imager and analyzed using the Quantity One software (version 4.4.0, Bio-Rad). Western blot data demonstrate successful biotinylation of IRF4 in *ex vivo* generated CD4$^{+}$ T cells from IRF4$^{Bio}$ mice. Moreover, data indicate comparable IRF4 expression levels in CD4$^{+}$ T cells derived from IRF4$^{Bio}$ and ROSA26$^{BirA}$ (ctrl) mice.

### Supplementary Fig. 7

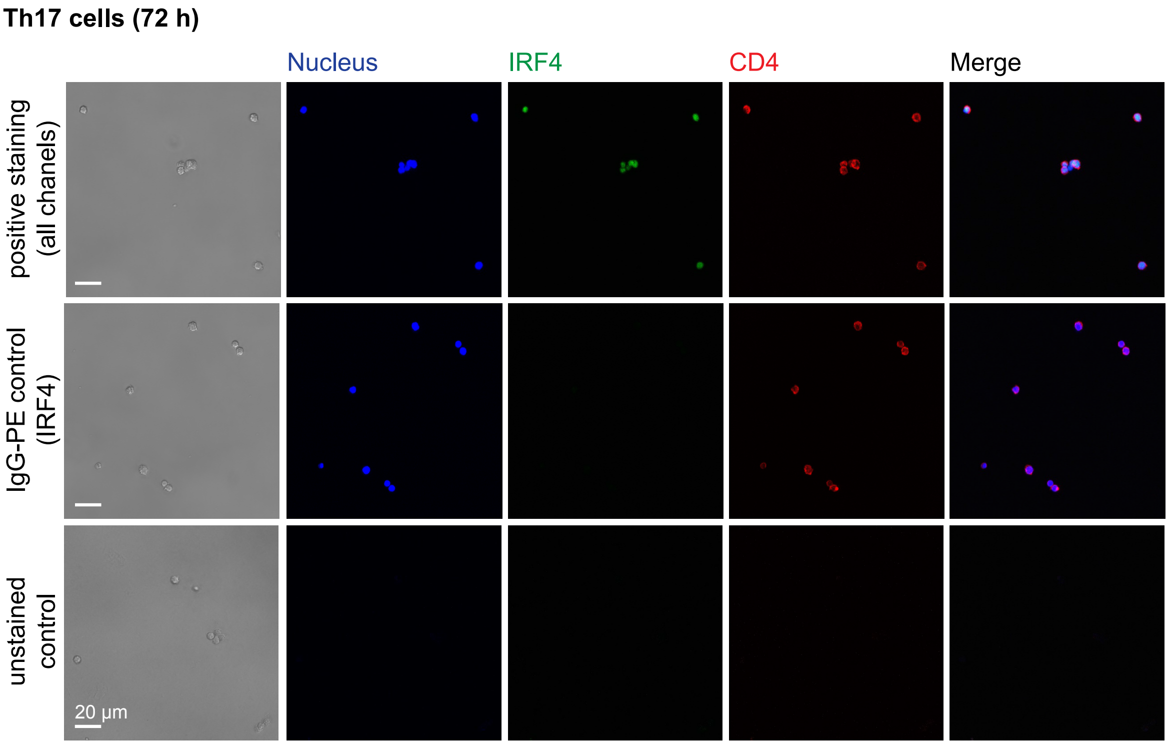


**Localization and expression of IRF4 three days after isolation of naïve CD4$^+$ T cells and Th17 *ex vivo* differentiation.** After three days of *ex vivo* differentiation, murine Th17 cells display high expression levels of IRF4 in the nucleus as indicated by immunofluorescence analysis (see also Fig. 2d). Cells were stained using phycoerythrin (PE)-conjugated anti-IRF4 as well as allophycocyanin (APC)-conjugated anti-CD4 antibodies (upper panel). IgG-PE served as negative control for the IRF4 staining (middle panel). Lower panel displays control with no stain. Scale bar 20 µm.

### Supplementary Fig. 8

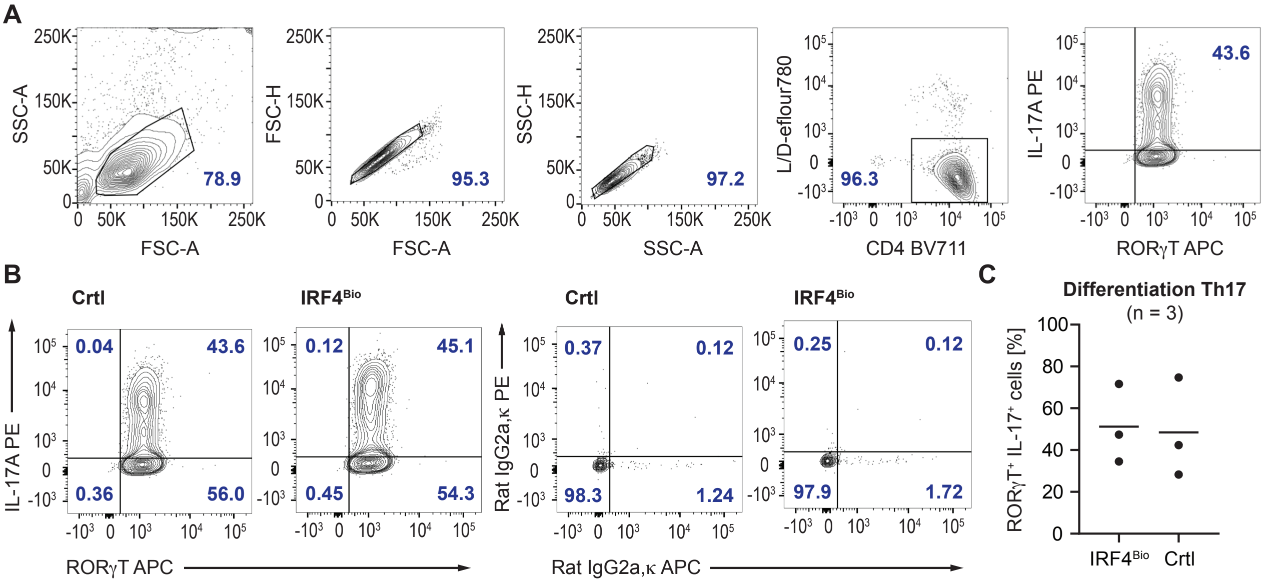


**Exemplified gating strategies for FACS analysis and characterization of murine Th17 cells**. (A) Gating strategy for the analysis of murine Th17 cells. (B) FACS analysis of Th17 cells three days after *ex vivo* differentiation of naïve $CD4^+$ T cells isolated from $IRF4^{Bio}$ and control mice ($ROSA26^{BirA}$). (C) Th17 cells used for the analyses of IRF4 interaction partners (three replicate experiments). Percentages of $ROR\gamma t^+$ $IL\text{-}17^+$ cells indicate successful *ex vivo* differentiation for $IRF4^{Bio}$ and control Th17 cells.

**Supplementary Fig. 9**

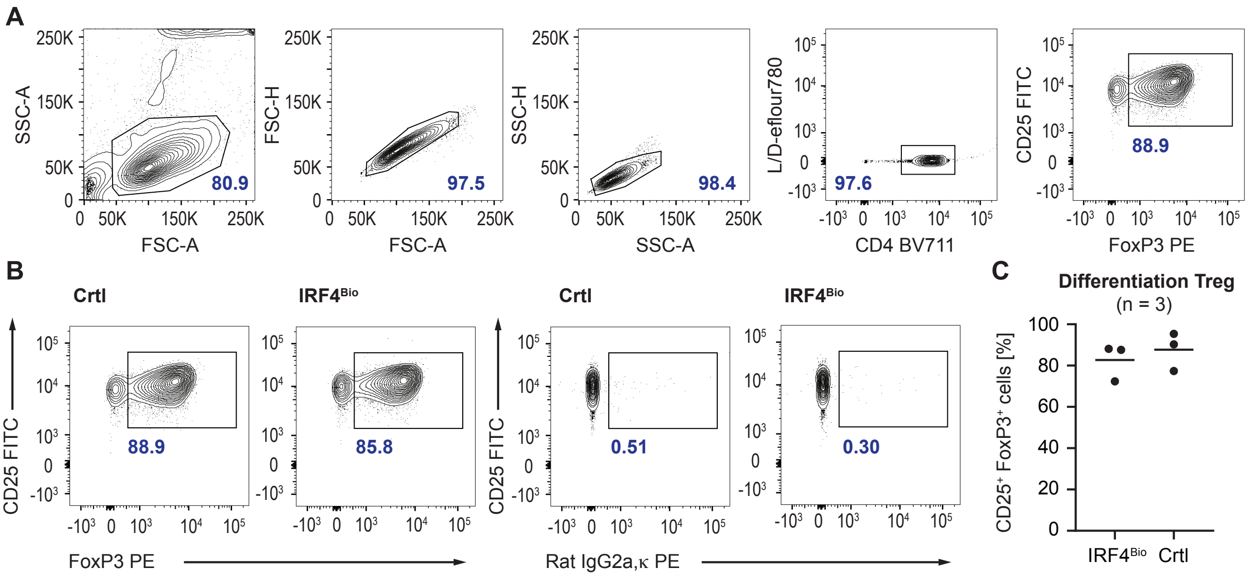


**Exemplified gating strategies for FACS analysis and characterization of murine iTreg cells**. (A) Gating strategy for the analysis of murine iTreg cells. (B) FACS analysis of iTreg cells three days after *ex vivo* differentiation of naïve $CD4^+$ T cells isolated from $IRF4^{Bio}$ and control mice ($ROSA26^{BirA}$). (C) iTreg cells used for the analyses of IRF4 interaction partners (three replicate experiments). Percentages of $CD25^+$ $FOXP3^+$ cells indicate successful *ex vivo* differentiation for $IRF4^{Bio}$ and control iTreg cells.

### Supplementary Fig. 10

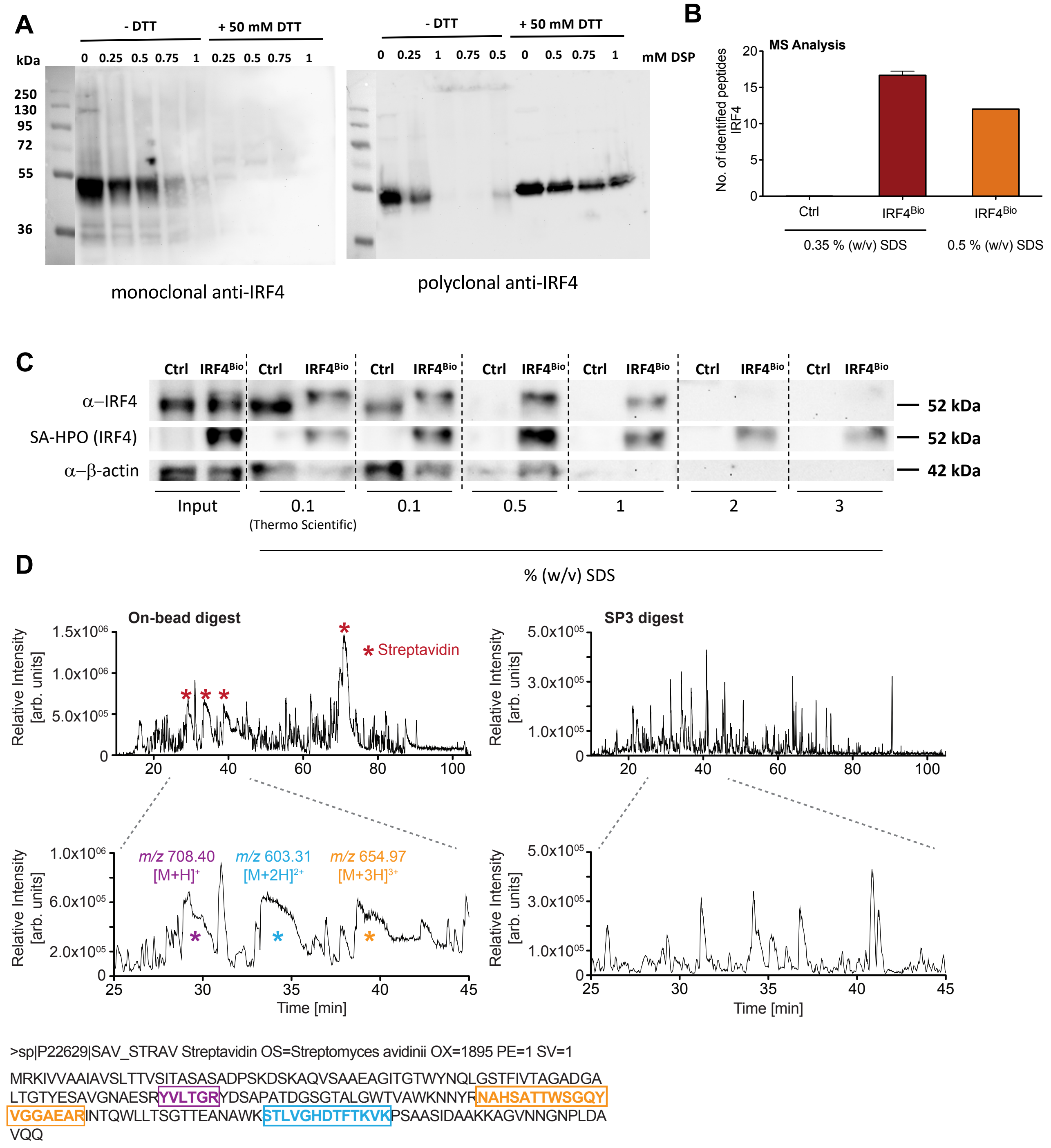


**Optimization of the IRF4$^{Bio}$ affinity purification (AP) protocol.** (A) Titration of Dithiobis[succinimidyl propionate] (DSP) for the intracellular cross-linking of IRF4 complexes. Western blot analysis of IRF4 in whole cell lysates of differentiated Th17 cells (time point: 72 h) treated with the indicated concentrations of DSP ranging from 0.25 mM to 1 mM DSP for 30 min at room temperature as described by Smith *et al.*[2]. For Western blot analysis, samples were prepared under non-reducing and reducing conditions as indicated. The present figure shows a dose-dependent reduction in monomeric IRF4. In case of the polyclonal antibody, a simultaneous appearance of cross-linked complexes across the top of the gel can be observed (right panel). Most of the monomeric IRF4 is decreased at concentrations between 0.5 – 0.75 mM and almost absent at 1.0 mM (absent in case of the immunoblot stained with the polyclonal antibody, less sensitive) indicating that the vast majority of IRF4 is cross-linked into high molecular complexes at these concentrations. In line with previous findings[2] monomeric IRF4 can be recovered by addition of DTT. Of note, cross-linked protein complexes could not be visualized with the monoclonal antibody, even after reduction, IRF4 was not detected (left panel). As the epitope recognized by the antibody contains lysine residues that are modified by crosslinker. Cross-linking in the final, optimized protocol was conducted using a concentration of 0.75 mM DSP. (B, C) Optimization of wash conditions for the isolation of IRF4 complexes. Nuclear lysates of *ex vivo* differentiated $CD4^+$ T cells from $IRF4^{Bio}$ and $ROSA26^{BirA}$ (Ctrl) mice were subjected to biotin-streptavidin affinity pulldown assays. Streptavidin-beads incubated with nuclear lysates were rinsed with RIPA buffer containing varying amounts of SDS. Eluates were analyzed by (B) mass spectrometry and (C) Western Blot using an anti-IRF4 antibody to detect IRF4 and horseradish peroxidase (HPO)–conjugated streptavidin identifying biotinylated IRF4. Proteins eluted from the streptavidin-beads after capture of biotinylated IRF4 indicate unspecific binding of unbiotinylated IRF4 using RIPA-buffer supplemented with 0.1 % (w/v) SDS as wash buffer. Using 0.5 % (w/v) SDS, nonspecifically bound IRF4 can be eliminated in the controls. However, higher amounts of SDS (above 0.5 % (w/v)) also lead to loss of biotinylated IRF4. We opted for an SDS concentration of 0.35 % (w/v) in the final protocol, as no unspecific binding of IRF4 was observed in the controls upon mass spectrometric analysis. Moreover, in triplicate MS analyses we detected a slightly higher number of proteotypic IRF4 peptides (16/17 unique peptides) using 0.35 % (w/v) SDS in the wash buffer as compared to a final concentration of 0.5 % (w/v) SDS (12 unique peptides). (D) On-bead digest versus SP3 digest. We were able to markedly reduce streptavidin contamination in the pull-down experiments (see upper and lower right panels) using SP3 for sample processing. Here, beads were initially boiled in a biotin-containing SDS-based buffer followed by SP3 digest[5,6], which results in a marked reduction of streptavidin contamination.

**Supplementary Fig. 11**

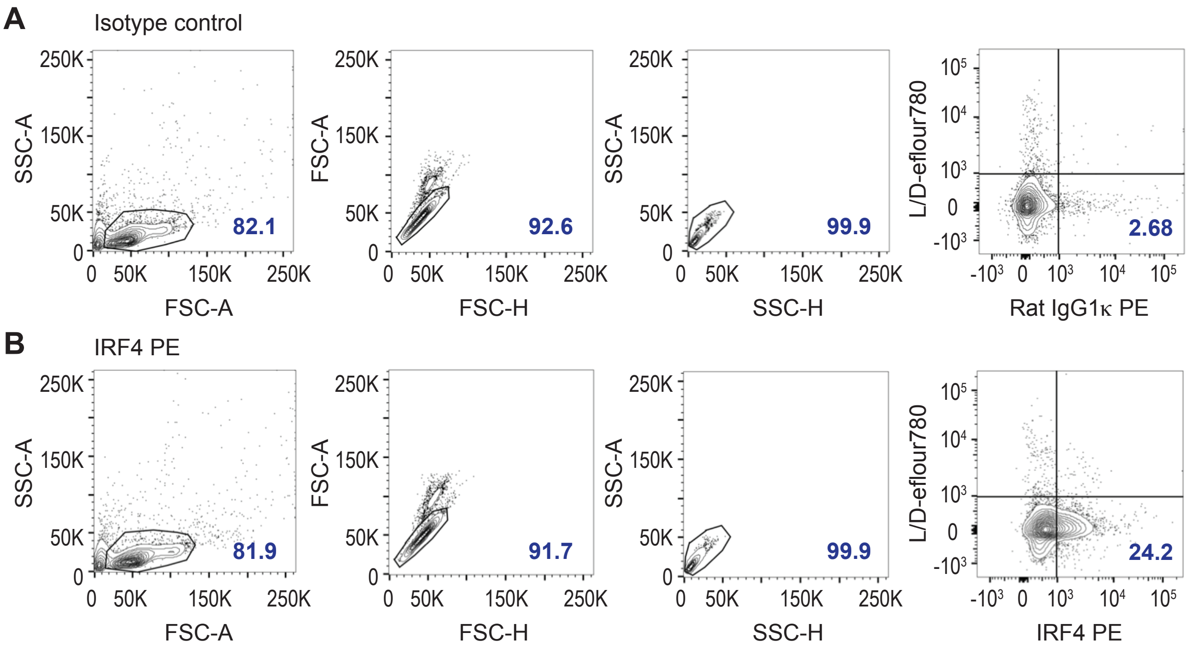


**FACS analysis of splenocytes and characterization of IRF4 positive subpopulations**. (A) Rat IgG1κ PE isotype control staining. (B) Around 24 % of the splenocytes used for initial streptavidin-biotin pulldown experiments (see also Fig. 2f) expressed IRF4.

**Supplementary Fig. 12**

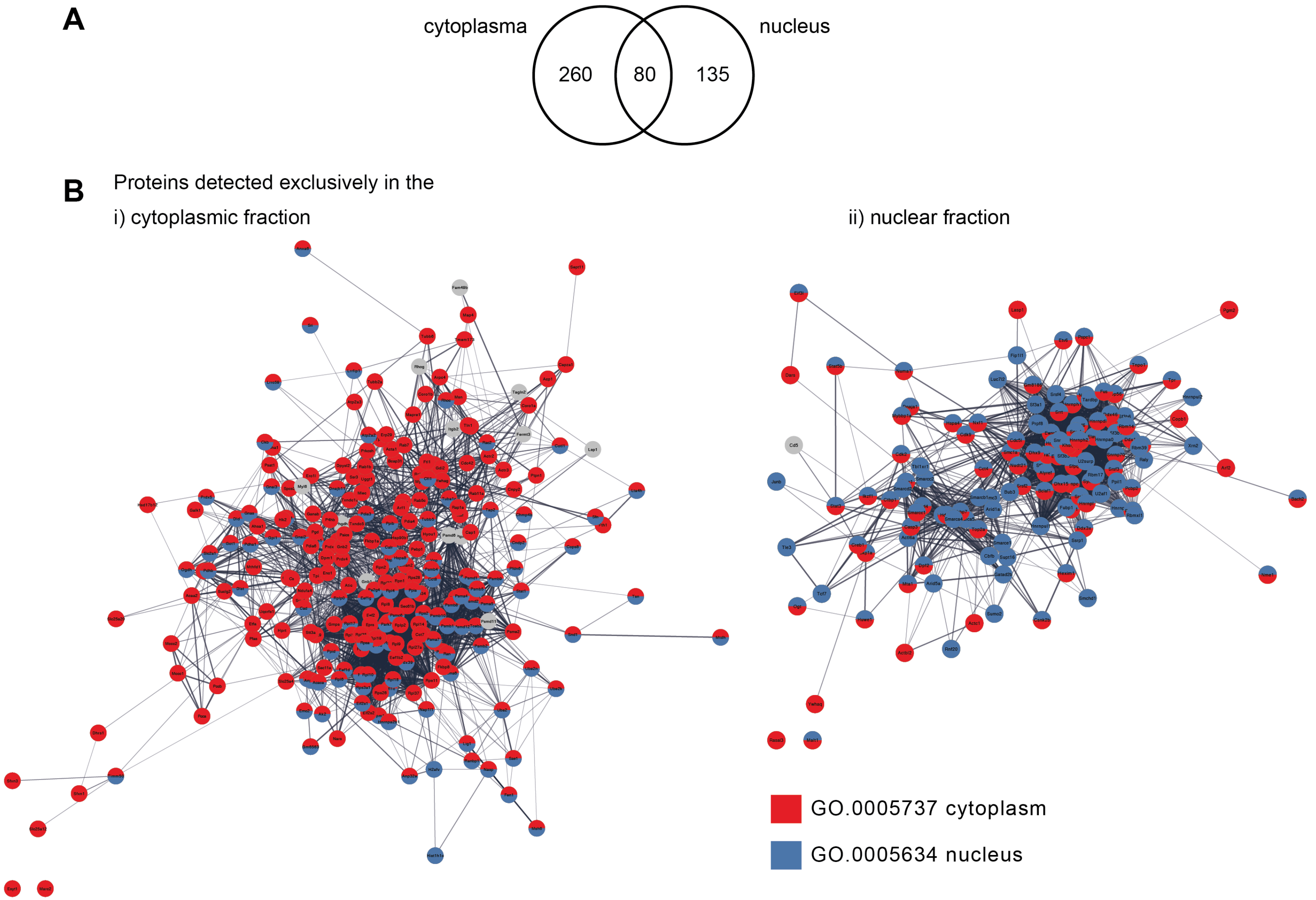


**Optimized AP-MS protocol allows the *ex vivo* characterization of transcription factor interactomes from different subcellular localizations.** Analysis of the IRF4 interactome in the nuclear and cytoplasmic fraction of *ex vivo* differentiated Th17 cells. Proteins exclusively detected in the cytoplasmic fraction are mainly associated with the cytoplasm and those in the nuclear fraction with the nucleus demonstrating the feasibility of our optimized protocol for the analysis of protein-protein interactions in different subcellular regions.

**Supplementary Fig. 13**

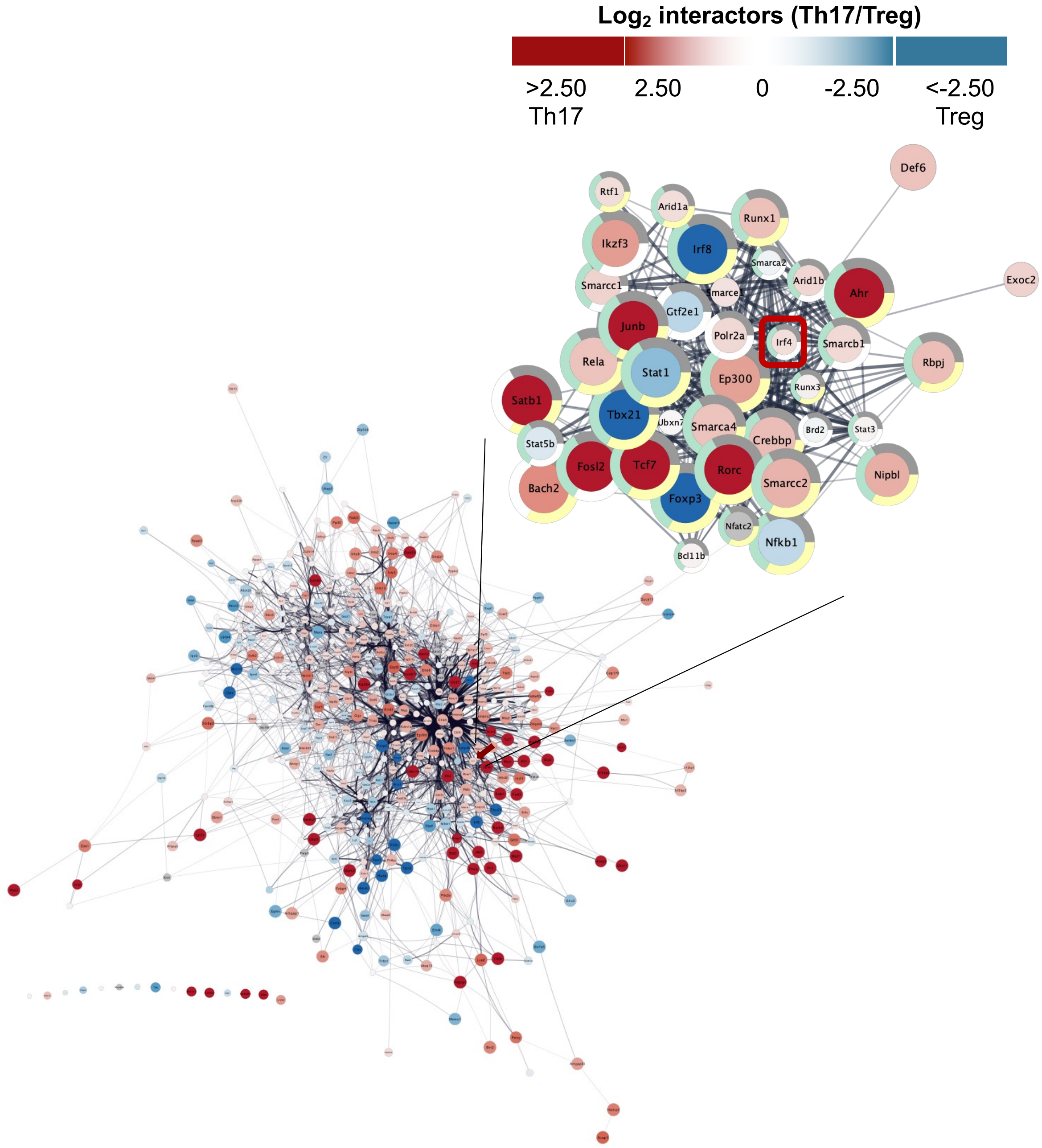


**Nuclear IRF4 interactome in fully differentiated Th17 and iTreg cells.** Full IRF4 protein interaction network in *ex vivo* generated Th17 and iTreg cells at day 3 as identified by AP-MS combining data from three independent AP experiments. In total, we identified 440 IRF4 interactors across both cell types. Functional network analysis of these IRF4 interplayers drawing upon the STRING database of protein-protein interactions indicated a strong network enrichment, where 422 out of the 440 candidates were assigned to a single interconnected network. STRING network analysis indicated 39 direct associations, either functional and/or physical, between IRF4 and the proteins detected by AP-MS (enlarged, also displayed in Figure 3a). Color code indicates enrichment either in Th17 (red) or iTreg cells (blue).

### Supplementary Fig. 14

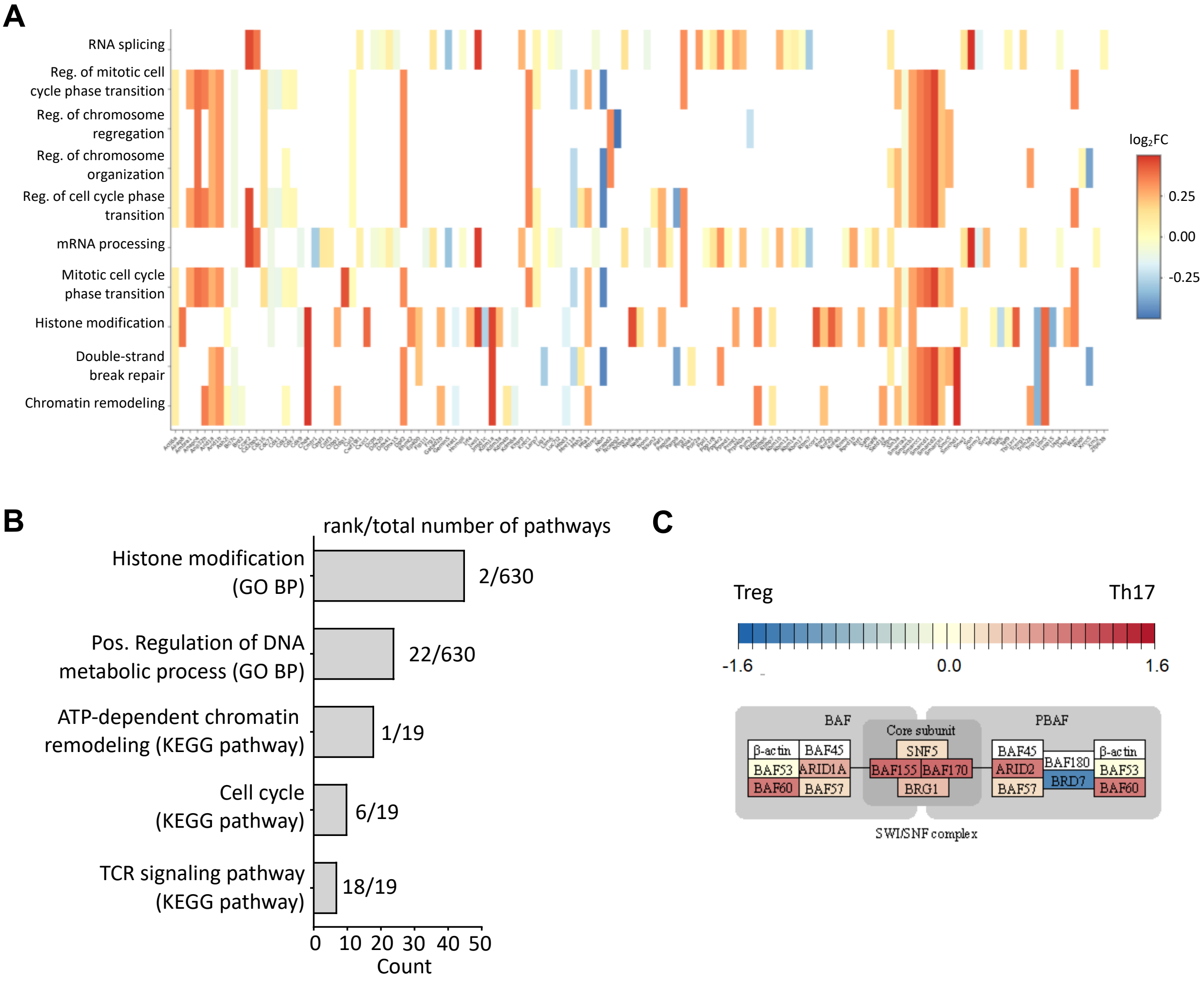


**Pathway analysis of proteins from the "IRF4 core" interactome reveals shared features between Th17 and iTreg cells.** (A) Heatmap of core interactome proteins that are mapped to the eight most enriched GO terms, including proteins associated with mRNA processing, histone modification and chromatin organization. (B) Number of proteins that are found in selected pathways (GO, KEGG, Reactome). The rank of the pathway is indicated on the right. (C) Many proteins from the SWI/SNF (switch/sucrose nonfermenting) and NuRD complexes (not displayed here) were identified in the core interactome, few were cell type specific. The NuRD complex, one of the major ATP-dependent chromatin remodeling complexes in cells, is crucial for the regulation of gene transcription, genome integrity as well as cell cycle progression acting broadly at many enhancers and promoters to dampen and fine-tune active gene expression[7]. As transcriptional activator, SWI-SNF has been reported to have an antagonistic role to NuRD on common regulatory elements competing for access to chromatin and generating opposite chromatin states[7]. SWI/SNF complexes can be divided into two broad classes depending on whether they contain one of two ATPases, BRG1 or hBRM. In our dataset, we detected the BRG1-BAF complex described as major regulator of the pluripotency of embryonic stem cells (hESCs)[8]. BRG1 plays a pivotal role in (STAT6-mediated) Th2 differentiation and Th2 cytokine transcription[9]. Upon IFNγ stimulation, phosphorylated STAT1 recruits BRG1 to IFNγ-activated sequences resulting in elevated gene transcription in primary T lymphocytes[10] suggesting a role of IRF4/STAT1-mediated regulation of BRG1 in our data set.

Colour code indicates enrichment in Th17 (red), iTreg cells (blue) or the core interactome (yellow).

**Supplementary Fig. 15**

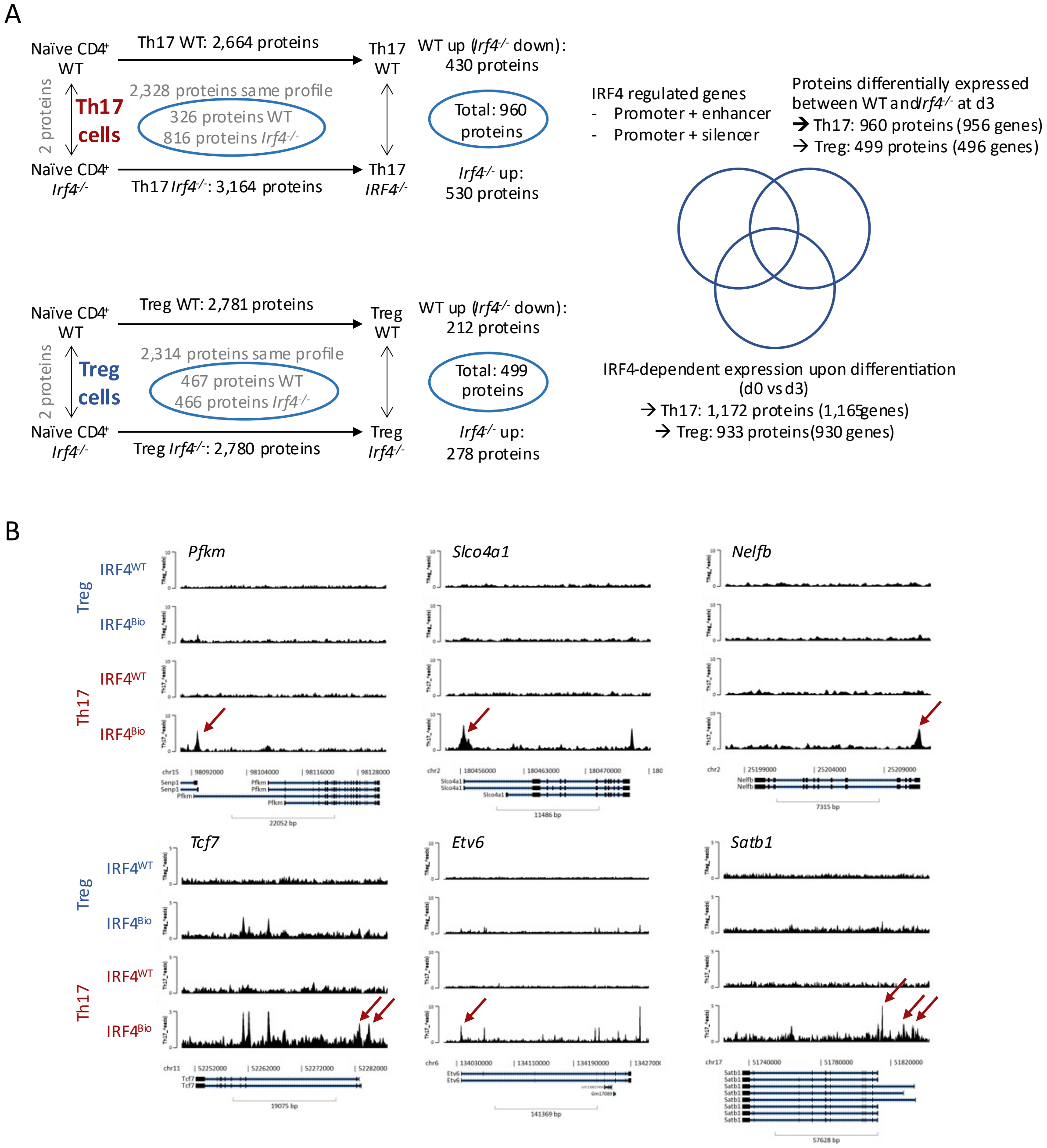


**Integrating proteome and ChIP-Seq analysis for the detection of proteins transcriptionally regulated by IRF4 which display aberrant expression levels in fully differentiated cells.** (A) Proteins that show aberrant expression patterns over time (i.e., d0 and d3) upon lack of IRF4 as well as differentially expressed proteins in *Ifr4*$^{-/-}$ Th17 and iTreg cells (at d3) were selected for further analysis: Overlap of selected proteins and IRF4 regulated genes (promotor region binding) as displayed in Fig. 6a and b. (B) ChIP-seq binding tracks for selected loci in iTreg and Th17 cells at 72 h. Red arrows indicate enhancer regions.

### Supplementary Fig. 16

**A**

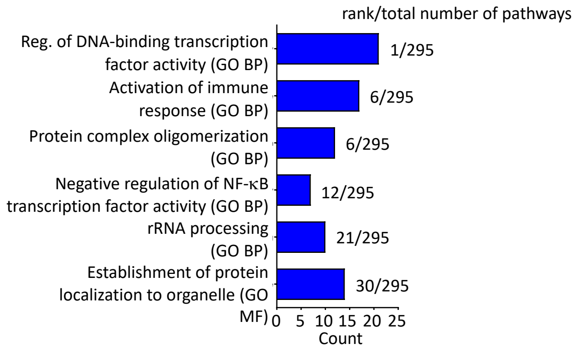


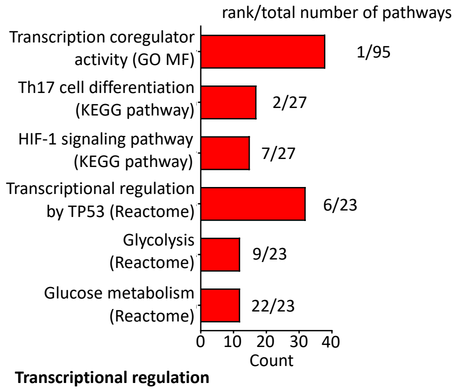


**B**

**Glycolysis**

Th17 cells

| | *Aldoa* | *Eno1* | *Foxk1* | *Gapdh* | *Gpi1* | *Pfkl* | *Pfkm* | *Pfkp* | *Pgk1* | *Pkm* | *Pgm1* | *Slc2a3* | *Hk3* |
|---|---|---|---|---|---|---|---|---|---|---|---|---|---|
| IRF4-BACH2 | nd | p + e | no p | nd | nd | nd | nd | nd | nd | nd | no p | no p | nd |
| IRF4-FLI1 | p | p + e | no p | p + e | p + e | nd | p + e | p + e | p + e | p + e | p + e | p + e | p + e |
| IRF4-GTF2I | nd | nd | no p | nd | nd | nd | nd | no p | nd | p + e | nd | no p | p |
| IRF4-AHR | p | p + e | no p | p + e | p + e | p + e | p + e | p + e | p + e | p + e | p + e | p + e | p + e |
| IRF4-EP300 | p | p + e | no p | p + e | p + e | p + e | p + e | p + e | p + e | p + e | p + e | p + e | p + e |
| IRF4-TCF7 | nd | p + e | no p | nd | p + e | nd | p + e | p + e | p + e | p + e | p + e | p + e | p + e |
| IRF4-RFX1 | p | p + e | no p | p + e | p + e | p + e | p + e | p + e | p + e | p + e | p + e | p + e | p + e |
| IRF4-SATB1 | nd | p + e | no p | p + e | p + e | p + e | p + e | p + e | nd | p + e | p + e | p + e | p + e |

Treg cells

| | *Aldoa* | *Eno1* | *Foxk1* | *Gapdh* | *Gpi1* | *Pfkl* | *Pfkm* | *Pfkp* | *Pgk1* | *Pkm* | *Pgm1* | *Slc2a3* | *Hk3* |
|---|---|---|---|---|---|---|---|---|---|---|---|---|---|
| IRF4-BACH2 | nd | nd | nd | nd | nd | nd | nd | nd | nd | nd | nd | nd | nd |
| IRF4-FLI1 | nd | nd | no p | nd | nd | nd | nd | no p | nd | p + e | nd | no p | nd |
| IRF4-FOXP3 | nd | no p | no p | p + e | nd | nd | no p | p + e | nd | p + e | no p | no p | nd |
| IRF4-GTF2I | nd | nd | no p | nd | nd | nd | nd | no p | nd | nd | nd | nd | nd |
| IRF4-EP300 | p | nd | no p | p + e | nd | nd | no p | p + e | nd | nd | no p | no p | nd |
| IRF4-TCF7 | nd | nd | no p | nd | nd | nd | nd | no p | nd | p + e | no p | no p | nd |
| IRF4-IRF8 | p | nd | nd | nd | nd | nd | no p | no p | nd | p + e | nd | no p | nd |
| IRF4-RFX1 | nd | no p | no p | p + e | nd | nd | no p | no p | nd | nd | no p | no p | nd |
| IRF4-SATB1 | nd | nd | nd | p + e | nd | nd | no p | p + e | nd | nd | no p | nd | nd |

**Transcriptional regulation**

Th17 cells

| | *Bach2* | *Ahr* | *Fli1* | *Gtf2i* | *Ep300* | *Tcf7* | *Satb1* | *Rfx1* | *Irf8* | *Foxp3* |
|---|---|---|---|---|---|---|---|---|---|---|
| IRF4-BACH2 | no p | no p | no p | p + e | nd | p + e | nd | nd | no p | nd |
| IRF4-FLI1 | no p | p + e | p + e | p + e | p + e | p + e | p + e | p + e | p + e | nd |
| IRF4-GTF2I | no p | p + e | no p | p + e | p + e | no p | p + e | p + e | p + e | nd |
| IRF4-AHR | no p | p + e | p + e | p + e | p + e | p + e | p + e | nd | p + e | nd |
| IRF4-EP300 | no p | p + e | p + e | p + e | p + e | p + e | p + e | p + e | p + e | nd |
| IRF4-TCF7 | no p | p + e | no p | p + e | p + e | p + e | p + e | p + e | p + e | nd |
| IRF4-RFX1 | no p | p + e | p + e | no p | p + e | p + e | p + e | p + e | p + e | nd |
| IRF4-SATB1 | no p | p + e | p + e | p + e | no p | p + e | p + e | p + e | p + e | nd |

Treg cells

| | *Bach2* | *Ahr* | *Fli1* | *Gtf2i* | *Ep300* | *Tcf7* | *Satb1* | *Rfx1* | *Irf8* | *Foxp3* |
|---|---|---|---|---|---|---|---|---|---|---|
| IRF4-BACH2 | no p | no p | nd | nd | nd | nd | nd | nd | nd | nd |
| IRF4-FLI1 | no p | no p | no p | no p | nd | no p | nd | nd | no p | nd |
| IRF4-FOXP3 | no p | no p | no p | no p | p + e | no p | no p | nd | no p | nd |
| IRF4-GTF2I | no p | nd | no p | no p | no p | nd | nd | nd | no p | nd |
| IRF4-EP300 | no p | no p | p + e | nd | p + e | no p | nd | nd | no p | nd |
| IRF4-TCF7 | no p | no p | no p | p + e | no p | no p | no p | nd | no p | nd |
| IRF4-IRF8 | no p | no p | no p | nd | no p | no p | no p | nd | no p | nd |
| IRF4-RFX1 | no p | no p | no p | no p | p + e | no p | nd | nd | no p | nd |
| IRF4-SATB1 | no p | no p | no p | no p | nd | no p | nd | nd | no p | nd |

**Pathway enrichment analysis of proteins that are regulated by IRF4 on DNA and protein level reveals cell type-specific functions.** (A) Number of proteins that are found in selected pathways (GO, KEGG, Reactome) of iTreg (blue, right panel) and Th17 cells (red, left panel). The rank of the pathway is indicated on the right. No enriched KEGG and Reactome pathways were detected in iTreg cells. (B) IRF4 DNA binding patterns in Th17 and iTreg cells. We analysed which of the genes targeted by the IRF4 complex contained composite motives indicating involvement of IRF4 binding partners in the regulation of the respective target genes. Particularly, in Th17 cells we found a lot of IRF4 interactors associated with transcriptional regulation that are involved in positive feedback loop regulation. The *Foxp3* gene is neither targeted in Th17 or iTreg cells. Here, no peaks were detected at all.

p + e: binding in promoter-enhancer region

p: binding in promoter region

no p: binding to target gene in distal intergenic or intron regions

n. d.: no binding detected at all, i.e. or no peaks detected in target gene (containing respective IRF4 composite motif)
